# Targeting tRNA-Arg-TCT-4-1 suppresses cancer cell growth and tumorigenesis

**DOI:** 10.64898/2026.02.19.706891

**Authors:** Esteban A. Orellana, Isobel E. Bowles, Xin Yang, Anais Torres, Sally R. Jamieson, Raja H. Ali, Alejandro Gutierrez, Richard I. Gregory

**Affiliations:** Department of Molecular and Systems Biology, Geisel School of Medicine at Dartmouth. Hanover, NH, 03755 USA; Dartmouth Cancer Center, Dartmouth. Lebanon, NH, 03756 USA; Division of Hematology/Oncology, Boston Children’s Hospital, Boston, MA 02115, USA; Department of Molecular, Cell, and Cancer Biology, UMass Chan Medical School, Worcester, MA 01605, USA; International Summer Research Experience (ISURE) at Dartmouth, Geisel School of Medicine at Dartmouth. Hanover, NH, 03655 USA; School of Arts and Sciences, Dartmouth College, Hanover, NH, 03755 USA; Medical Genomics Research Department, King Abdullah International Medical Research Center (KAIMRC), King Saud Bin Abdulaziz University for Health Sciences (KSAU-HS), King Abdulaziz Medical City (KAMC), Ministry of National Guard Health Affairs (MNG-HA), Riyadh, Saudi Arabia; Department of Oncology, St. Jude Children’s Research Hospital, Memphis, TN 38105, USA

**Keywords:** tRNA, tRNA-Arg-TCT-4-1, isodecoder, mRNA translation, antisense, cancer, inhibition

## Abstract

tRNAs play a critical role in protein synthesis, influencing mRNA translation dynamics to shape proteomes. Emerging evidence links dysregulated tRNA activity to cancer progression, with tRNA-Arg-TCT identified as an oncogenic driver when ectopically overexpressed in non-malignant cells. The requirement of endogenous tRNA-Arg-TCT in cancer biology, however, remains untested. Moreover, considering that the tRNA-Arg-TCT family comprises six genes in humans, the importance of an individual tRNA isodecoder in cancer remains unknown. Here, we find elevated levels of tRNA-Arg-TCT-4-1 isodecoder are associated with poor patient prognosis across multiple cancer types. We demonstrate that, using different antisense RNA strategies, specific inhibition of tRNA-Arg-TCT-4-1 suppresses the growth of glioblastoma (GBM) and liposarcoma (LPS) cancer cells. Mechanistically, we find that tRNA-Arg-TCT-4-1 inhibition leads to a codon-biased remodeling of mRNA translation and the proteome, preferentially suppressing expression of growth-promoting genes and pathways encoded by mRNAs enriched in arginine AGA codons. Strikingly, intratumoral delivery of an antisense oligonucleotide (ASO) targeting tRNA-Arg-TCT-4-1 suppresses tumor growth and extends survival in mouse xenograft experiments performed using either a human LPS cell line or a patient-derived soft tissue sarcoma model. This study provides a foundation for targeting tRNA dysregulation as a novel therapeutic approach for cancer.

**One Sentence Summary:** This study identifies tRNA-Arg-TCT-4-1 as a new anti-cancer therapeutic target and demonstrates that an antisense oligonucleotide (ASO) targeting this tRNA effectively suppresses tumorigenesis and extends survival in mouse xenograft models.

## Introduction

Transfer RNAs (tRNAs) play a crucial role in protein synthesis, influencing the dynamics of messenger RNA (mRNA) translation(*1*). Historically, tRNAs have been regarded as housekeeping molecules, lacking specific regulation due to their high cellular abundance and stability. However, recent studies (*2–9*), including our own work(*1*, *10*), have revealed the opposite: tRNA levels are tightly regulated, and even subtle changes in abundance or post-transcriptional modifications profoundly affect translation and protein levels, resulting in disease (*1*). While much research has focused on the regulation of gene expression through transcription and mRNA synthesis, comparatively little is known about how aberrant translation contributes to a defective proteome. tRNA dysregulation, through alterations in the abundance or modification status of specific tRNAs, has only recently been recognized as a significant contributor to metabolic disorders, neurological diseases, and cancers(*1*).

MicroRNA (miRNA) sequencing data repurposed from The Cancer Genome Atlas (TCGA) has been utilized to assess the variations in tRNA levels between diseased and healthy tissues. The findings reveal that tRNA dysregulation is a prevalent occurrence across various human cancers (*10–32*)This observation affects various human cancers since increasing evidence indicates that changes in the tRNA pool leads to codon-biased reprogramming of mRNA translation, which can specifically promote oncogenic activity instead of globally impacting protein synthesis programs. Translation initiation represents the initial rate-limiting step in protein synthesis and can be exploited to promote tumorigenesis (*33*, *34*). For instance, increased levels of tRNA-iMet lead to alterations in the metabolic and growth rates of immortalized human breast cells(*32*); this overexpression also encourages metastasis in melanoma tumors(*22*) and significantly elevates tumor burden and vascularization mice(*22*). An additional regulatory layer in the translation of oncogenic mRNAs involves modifications to elongator tRNAs, leading to changes in cognate codon usage. The overexpression of tRNA-Glu-UUC and tRNA-Arg-CCG fosters a pro-metastatic state in breast cancer by increasing the translation of transcripts that are rich in their corresponding codons(*13*). Moreover, a genome-wide loss-of-function CRISPR–Cas9 screen revealed tRNA-Val as a key adaptation in the pathobiology of leukemia(*15*).

We and others have demonstrated that the upregulation of the N7-methylguanosine (m^7^G) methyltransferase (MTase), METTL1, results in enhanced m^7^G tRNA modification and stability, ultimately reshaping the tRNA pool(*10*, *35*). METTL1/WDR4 is responsible for depositing a methylation mark on guanine 46 (m^7^G46) in a subset of tRNAs, which stabilizes the tertiary folding of tRNAs and contributes to increased stability(*36*). Elevated levels of METTL1/WDR4 result in the accumulation of specific tRNAs thereby changing the composition of the proteome by dictating which proteins are synthesized more efficiently in a codon-biased manner. The human genome encodes multiple tRNA isoforms. Remarkably, humans have over 500 tRNA genes, approximately 50% of which are isodecoders, which are isoforms that share identical anticodon sequences but otherwise diverge in their body sequence from related tRNAs. Isodecoders were long considered to be functionally redundant copies, providing a buffer against deleterious mutations in cognate tRNAs. However, recent evidence, including ours, challenges this assumption, suggesting that tRNA isodecoders may have distinct, specialized roles(*4*, *10*, *37*, *38*). For example, tRNA-Arg-TCT-4-1 is an isodecoder that, when overexpressed, is sufficient to phenocopy METTL1-mediated oncogenic transformation via reprogramming of codon-biased translation(*10*). Mechanistically, we have shown that the increased abundance of tRNA-Arg-TCT-4-1 in cancer results in the preferential translation of proteins that stimulate growth and are enriched with the respective cognate AGA codon(*10*)

tRNA-Arg-TCT-4-1 is part of a tRNA family with five individual members expressed from six genomic loci, and its normal expression is highly restricted to the central nervous system (CNS)(*37*). The proposed normal physiological role of tRNA-Arg-TCT-4-1 in the adult mouse brain is to regulate synaptic transmission (*38*). Interestingly, data from small RNA sequencing experiments from The Cancer Genome Atlas (TCGA) indicate that tRNA-Arg-TCT is aberrantly elevated in multiple cancer types, including breast, lung, and soft tissue sarcomas(*10*). Research has demonstrated that the DNA region containing tRNA-Arg-TCT-4-1 is significantly hypomethylated in cancers when compared to matched normal tissues (*39*). This indicates that, in addition to posttranscriptional mechanisms, epigenetic pathways may also regulate its abnormal expression in malignant contexts.

Even though altered tRNA activity can drive pathological processes in a codon-dependent manner, these mechanisms, to our knowledge, have not been explored in the context of therapeutic vulnerabilities of cancer. Therefore, given the oncogenic potential of tRNA-Arg-TCT-4-1 and its specificity for normal CNS tissue, which is protected by the blood-brain barrier, we asked whether this tRNA isodecoder could be a dependency for cancer. Here, we show that specific inhibition of tRNA-Arg-TCT-4-1 is feasible and abolishes tumor formation *in vivo*. Mechanistically, we show that tRNA-Arg-TCT-4-1 alters the translation efficiency of genes involved in cell cycle regulation, increases ribosome pauses at AGA codons, and leads to corresponding changes in protein expression. We furthermore show the therapeutic efficacy of an antisense oligonucleotide (ASO) delivered to tumors in mouse xenograft models, which strongly inhibits tumor growth and substantially extends survival. Altogether, our data support the use of tRNA modulation strategies in precision medicine.

## Results

### tRNA-Arg-TCT-4-1 levels are elevated in human tumors and are associated with poor patient prognosis

Data mining from small RNA sequencing experiments from The Cancer Genome Atlas (TCGA) indicates that tRNA-Arg-TCT-4-1 is aberrantly expressed in multiple cancer types. Within datasets that include normal matched tissues, we found that tRNA-Arg-TCT-4-1 levels are significantly increased more than 2-fold in 13 cancer types with breast invasive adenocarcinoma (BRCA) the type with the highest and most significant upregulation (**Fig. 1A**), and its levels account for ∼1 to 10% of the tRNA-TCT pool isodecoder family (**Fig. S1A**). Interestingly, tRNA-Arg-TCT-4-1 is the most upregulated tRNA in BRCA (**Fig. 1B**) and is particularly enriched in the basal, Her2, LumA, and LumB molecular subtypes, which are generally regarded as aggressive (*40*), compared to a normal-like subtype (**Fig. 1C**). Notably, no significant enrichment was found based on disease stage (**Fig. S1B**). We then validated this observation using a breast cancer tissue microarray (TMA) with a modified tRNA-fluorescent in situ hybridization (FISH) protocol based on BaseScope technology (ACDBio). The specificity of the tRNA-Arg-TCT-4-1 probes was first confirmed using mouse brain tissue, compared to negative control probes and a tRNA-Phe-GAA probe as a positive control. This test showed that tRNA-Arg-TCT-4-1 expression is limited to neurons (**Fig. S1C**), which aligns with previous findings(*38*). Our TMA results confirmed the TCGA RNA sequencing findings, showing a markedly increased level of tRNA-Arg-TCT-4-1 localized in epithelial cancer cells, but not in the cancer stroma or normal epithelia in adjacent tissues (**Fig. 1D, E**). Additionally, elevated levels of tRNA-Arg-TCT-4-1 are negatively correlated with patient survival in at least 7 cancer types, with head and neck squamous cell carcinoma (HNSCC) showing the most significant difference (**Fig. 1F, S1D**). Given that not all cancer datasets in TCGA include normal-tissue control samples, we aimed to determine the relative expression levels of tRNA-Arg-TCT-4-1 across 24 cancer types. The data indicate that the cancer types with the highest levels of tRNA-Arg-TCT-4-1 are gliomas (LGG), paraganglioma (PCPG), followed by mesothelioma (MESO), and soft tissue sarcomas (SARC) (**Fig. 1G**). We also confirmed the increased level of tRNA-Arg-TCT-4-1 in SARC (liposarcomas) using tRNA-FISH. We again found that the high level of the isodecoder is primarily present in cancer cells, but not in benign tumors or normal tissues (**Fig. S1E, F**). Overall, the data suggest that the elevation of tRNA-Arg-TCT-4-1 levels is a widespread phenomenon in different human cancer types, and this upregulation is associated with a lower likelihood of patient survival.

**Figure 1.**
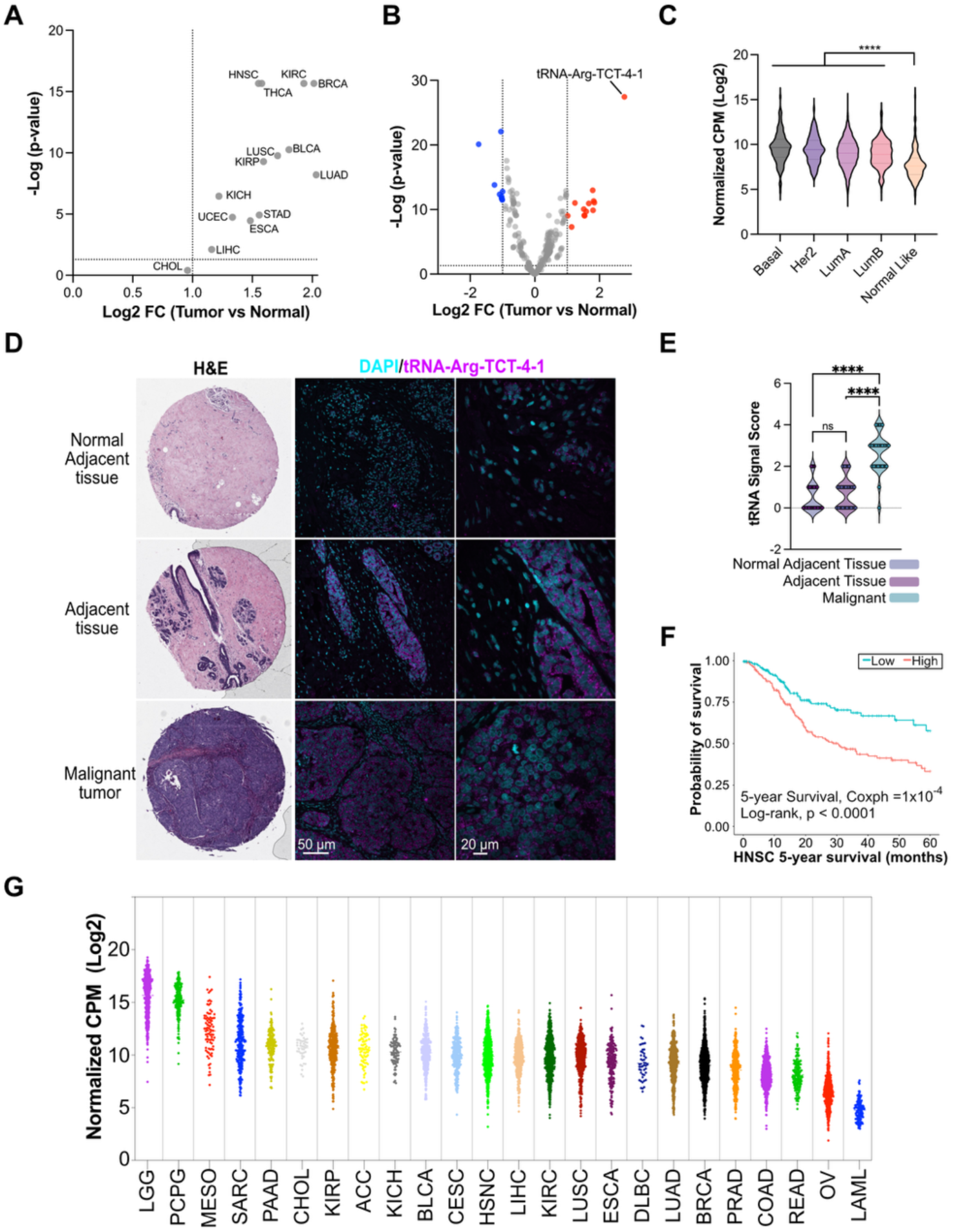
tRNA-Arg-TCT-4-1 is dysregulated in human cancers and associated with poor patient prognosis. **A)** tRNA-Arg-TCT-4-1 fold change (FC) expression between tumor vs normal samples. **B)** tRNA abundance in breast invasive carcinoma (BRCA). Data from TCGA. **C)** tRNA-Arg-TCT-4-1 abundance in various molecular subtypes of BRCA. CPM: counts per million. **D)** tRNA-Arg-TCT-4-1 expression in BRCA tumor samples using tRNA-FISH (BaseScope). **E**) Signal quantification from (D), full TMA. Error bars: Mean ± S.D. P values from Two-way ANOVA with posthoc Bonferroni correction. ns: not significant; ****p < 0.0001. **F**) Kaplan-Meier survival curve of HNSCC patients with low versus high tRNA-Arg-TCT-4-1 expression levels. Mean cut-off. Data are from TCGA. Wilcoxon and log-rank test. **G)** Relative tRNA-Arg-TCT-4-1 normalized levels in 24 human cancers. Data are from TCGA.

### Inhibition of tRNA-Arg-TCT-4-1 suppresses tumorigenesis

Considering that tRNA-Arg-TCT-4-1 overexpression is oncogenic in various contexts(*10*), we next sought to ask if tRNA-Arg-TCT-4-1 could be a possible therapeutic target. Considering a previous study utilized short hairpin RNA (shRNA) to deplete tRNA(*13*, *41*, *42*), we designed a shRNA to specifically knock down tRNA-Arg-TCT-4-1 in LNZ308 human glioma cells (**Fig. 2A**). Due to the small target sequence (73 nt), we were restricted to a single shRNA sequence that would target isodecoder 4-1 on the D-arm. In this cell line, the abundance of tRNA-Arg-TCT-4-1 constitutes only 0.88% of the total pool of tRNA-Arg-TCT, and the most abundant isodecoder is tRNA-Arg-TCT-1-1 with 53.74% of the total sequence read counts (**Fig. S2A**). We generated stable cell lines expressing either the control (shGFP) or the shRNA targeting tRNA-Arg-TCT-4-1 (shArg) in LNZ308 cells. To our surprise, shRNA-mediated targeting of tRNA-Arg-TCT-4-1 (shArg) did not reduce tRNA expression but instead led to a slight increase in tRNA-Arg-TCT-4-1 levels compared to the shGFP control (**Fig. 2B**). Moreover, tRNA sequencing analysis revealed that the observed increase in tRNA-Arg-TCT-4-1 is specific since only expression of this targeted tRNA was altered in the shArg cells (**Fig. 2C**). We postulated, therefore that shArg might still specifically inhibit tRNA-Arg-TCT-4-1 function, albeit by a possible sequestration (sponging) by the antisense shRNA rather than by a classical RNA-interference (RNAi) mechanism. To test this, we deployed a translation reporter assay that is sensitive to changes in active tRNA-Arg-TCT levels. This assay uses a fusion mCherry-Hmga2 construct enriched with AGA codons (Wt) and expresses an acGFP1 as a translation normalizer. A mutant sensor (Mut) without AGA codons is employed to assess the specificity of the changes. The data indicate that cells expressing the shArg construct have a significant decrease in the synthesis of the AGA-enriched sensor mCherry-Hmga2 compared to parental cells and the shGFP control, supporting a specific tRNA-Arg-TCT-4-1 loss of function by the shArg, and suggesting a potential steric hindrance mechanism that sequesters tRNA-Arg-TCT-4-1 molecules acting as a sponge to inhibit tRNA function (**Fig. 2D**). Consistent with this notion, we found no change in tRNA-Arg-TCT-4-1 aminoacylation in the shArg conditions (**Fig. S2B**). Next, we explored the phenotypic implications of tRNA inhibition by conducting soft agar colony formation assays to assess its effect on oncogenic cell growth. We found that tRNA-Arg-TCT-4-1 inhibition strongly reduces the ability of LNZ308 glioblastoma cells to form colonies (**Fig. 2E, F**). This result indicated that the ability of these cells to form tumors might be compromised; thus, we used a xenograft approach in immunocompromised mice to monitor tumor formation *in vivo*. Strikingly, tRNA-Arg-TCT-4-1 inhibition completely abolished the ability of these cancer cells to form tumors *in vivo* (**Fig. 2G, H**).

**Figure 2.**
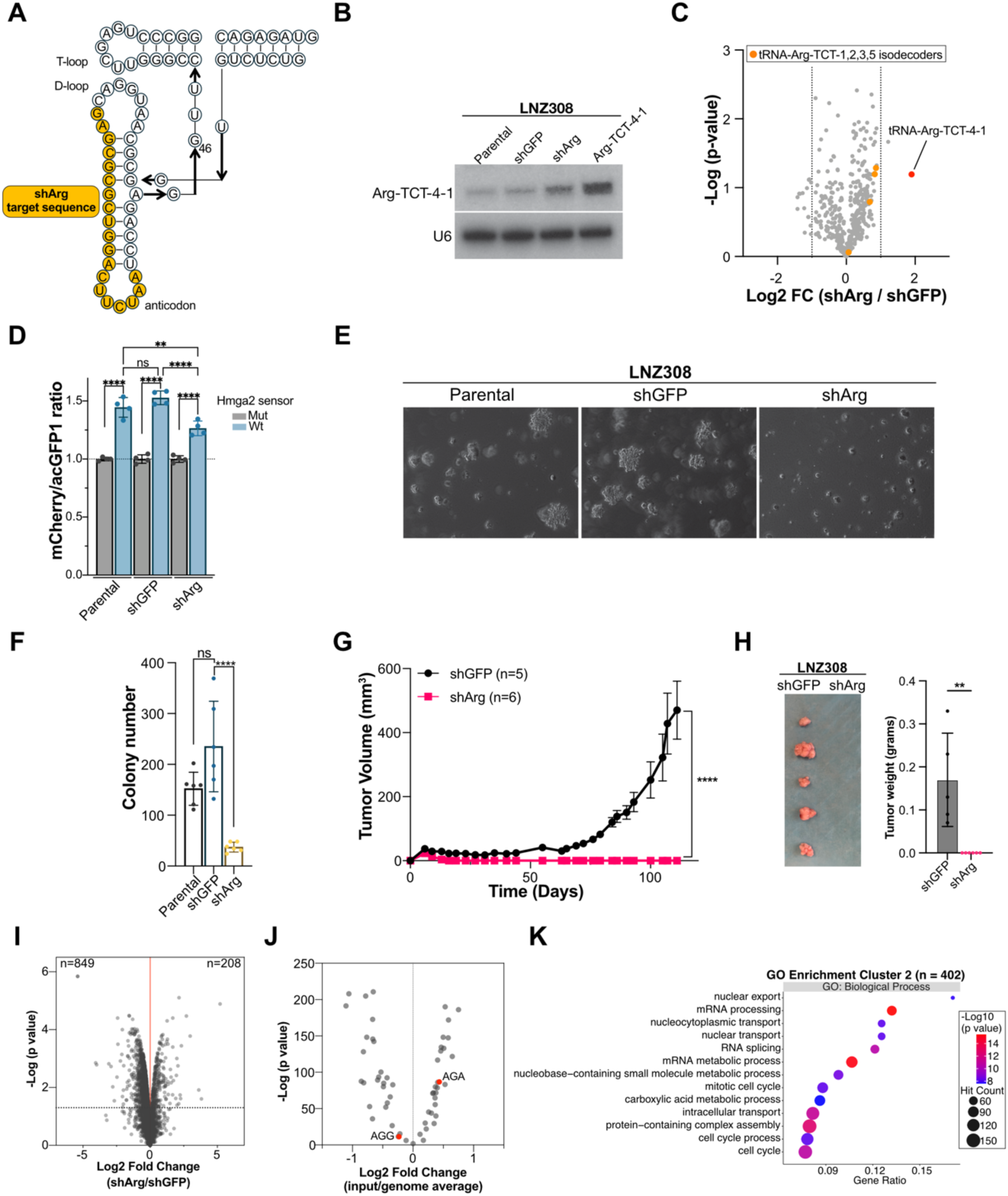
Inhibition of tRNA-Arg-TCT-4-1 prevents tumor formation in a glioblastoma model. **A)** Schematic of shArg targeting sequence on tRNA-Arg-TCT-4-1. **B)** Northern blot showing relative tRNA-Arg-TCT-4-1 levels. Arg-TCT-4-1 lane is an overexpression construct used as a control. **C)** Changes in tRNA abundance upon shArg treatment. Orange circles represent tRNA-Arg-TCT isodecoders 1,2,3 and 5. **D)** mCherry/acGFP1 ratios measured by flow cytometry between mCherry-Hmga2-WT (5 of 12 AGA codons) and mCherry-Hmga2-MUT (0 of 12 AGA codons) reporters. Data are mean ± SD. n = 2. p values from two-way ANOVA with Šídák correction. ^∗^p < 0.05 and ^∗∗^p < 0.01. **E)** Images of 3D soft agar colony formation assays in LNZ308 cells treated with shGFP or shArg. **F**) Quantification of colonies from (E). n=5, Error bars: Mean ± S.D. P values from Two-way ANOVA with posthoc Bonferroni correction. ns: not significant; ****p < 0.0001. **G)** *In vivo* tumor formation of LNZ308 cells (n = 5 or 6; error bars denote mean ± SEM). ^∗∗∗∗^p < 0.0001. Two-way analysis of variance (ANOVA) and Bonferroni correction. **H)** Tumor weight from (G). Error bars denote mean ± SD. p value from paired Student’s t test. **I)** Changes in protein abundance between shArg (heavy) expressing cells and shGFP (light) control cells measured by SILAC-based proteomics; n = 3, moderated t-test. **J)** Codon enrichment (gene input/global genome average) of genes in cluster 2 (from Fig. S2C). P-value from Fisher’s Exact Test with Benjamini–Hochberg FDR correction. **K)** Gene Ontology enrichment analysis using downregulated proteins in cluster 2 (from Fig. S2C) upon tRNA-Arg-TCT-4-1 inhibition. p<0.05, FC≤1.2.

We next sought to understand how the inhibition of tRNA-Arg-TCT-4-1 alters the proteome in these human glioblastoma cells. We performed stable isotope labeling by amino acids in cell culture (SILAC) proteomics comparing shArg versus shGFP cells (**Table S1**). The results show that inhibiting tRNA-Arg-TCT-4-1 leads to a broad dysregulation of the proteome in shArg-treated cells compared to the shGFP control, with most changes decreasing protein output with 849 genes significantly downregulated (<1.2-fold; p<0.05) vs 208 genes significantly upregulated (>1.2-fold; p<0.05) (**Fig. 2I**). Next, we considered that it was improbable that all the downregulated proteins resulted directly from tRNA inhibition, and that the list likely includes indirect effects. Therefore, we performed codon-enrichment analyses for each gene, compared them with the genome-wide average, and conducted hierarchical clustering. This led to the segregation of the gene list (736 genes with annotations) into gene clusters (rows) and codons (columns) (**Fig. S2C**). Interestingly, we observed a bias in how codons cluster by the third nucleotide, particularly for codons ending in G/C or A/T. Similarly, two major gene clusters emerged based on their enrichment with codons ending in A/T (i.e. AGA) or codons ending in C/G (**Fig. S2C**). This structure disappears if a randomly generated gene list of similar size is used for analysis (**Fig. S2D**). Gene cluster 2 showed the most substantial enrichment in AGA codons (**Fig. 2J**), while cluster 1 showed no enrichment (**Fig. S2E**). Functionally, gene cluster 2, enriched for AGA, is associated with genes involved in the cell cycle, mRNA processing/splicing, and intracellular transport (**Fig. 2K**). In contrast, cluster 1 is enriched for genes involved in nucleoside metabolism and various metabolic processes (**Fig. S2F**). Altogether, our data indicate that it is feasible to specifically inhibit tRNA-Arg-TCT-4-1, and that inhibiting this tRNA isodecoder leads to decreased expression of growth-promoting proteins and suppression of tumor growth *in vivo*.

### Inhibition of Arg-TCT-4-1 causes changes in mRNA translation and ribosome stalling

Since tRNA-Arg-TCT-4-1 is normally only expressed in the central nervous system (CNS)(*4*, *37*, *38*, *43*) and the possibility that tRNA-Arg-TCT-4-1 inhibition might only be effective in a CNS context, we next sought to determine whether our inhibition strategy would also be effective in other cancer types that are not part of the CNS. To this end, we selected sarcoma, specifically liposarcoma (LPS), because it exhibits among the highest levels of tRNA-Arg-TCT-4-1 expression (**Fig. 1G, S1A, S1E, S1F**). We first identified the LPS cell line LPS853, which has a moderate expression level of tRNA-Arg-TCT-4-1, and LPS141, which shows no detectable expression of the isodecoder, in order to generate stable cell lines expressing shArg (**Fig. 3A**). Based on the expression of tRNA-Arg-TCT-4-1, we hypothesized that if our shRNA specifically inhibits tRNA-Arg-TCT-4-1, only LPS853 cells would respond to the tRNA inhibition. Therefore, we first measured cell proliferation in a colony formation assay and found that shArg causes a significant decrease in LPS853 colonies compared to parental and shGFP controls; however, no significant difference was observed in LPS141 (**Fig. 3B**). Similar to our previous observation in LNZ308, shArg expression leads to a modest accumulation of the tRNA without changes in the aminoacylation level (**Fig. S3A**). We then performed LPS853 xenograft experiments in immunocompromised mice and found that shArg also prevents tumor formation *in vivo* (**Fig. 3C, S3B**). We utilized Ribo-Seq to investigate the impact of tRNA inhibition on mRNA translation and discovered that cells expressing shArg exhibit defects in translation efficiencies (TE), with 521 genes showing decreased TE (<1.5-fold, p<0.05) and 498 genes demonstrating increased TE (>1.5-fold, p<0.05) (**Fig. 3D; Table S2**). After discovering changes in translation efficiencies, we next investigated whether shArg-treated cells exhibit ribosome translocation defects (i.e., stalling), as indicated by ribosome pause scores. The results revealed a specific and significant increase in AGA pause score when tRNA-Arg-TCT-4-1 is inhibited (**Fig. 3E**). We reasoned that similar to the patterns observed in LNZ308 proteomic changes, the changes in TE would include both direct and indirect effects of the tRNA inhibition. Therefore, we conducted codon enrichment analyses again for each gene as previously described, and this resulted in the segregation of the gene list (521 annotated genes) into gene clusters (rows) and codons (columns), as shown in **Fig. 3F**. The clustering uncovered a bias in how codons are grouped based on their third nucleotide, particularly for those ending in G/C or A/T. Similarly, two primary gene clusters were identified, distinguished by enrichment for codons ending in A/T (e.g., AGA) or C/G (**Fig. 3F**). Gene cluster 1 showed the most substantial enrichment in AGA codons (**Fig. 3G**), while cluster 2 showed AGA depletion (**Fig. 3H**). Functionally, gene cluster 1, enriched for AGA, is associated with genes involved in chromatin remodeling, mRNA decay, mitochondrial autophagy, and neoplastic disease (**Fig. 3I**). In contrast, cluster 2, which is depleted for AGA, is primarily enriched for genes involved in neuron development (**Fig. 3J**).

**Figure 3.**
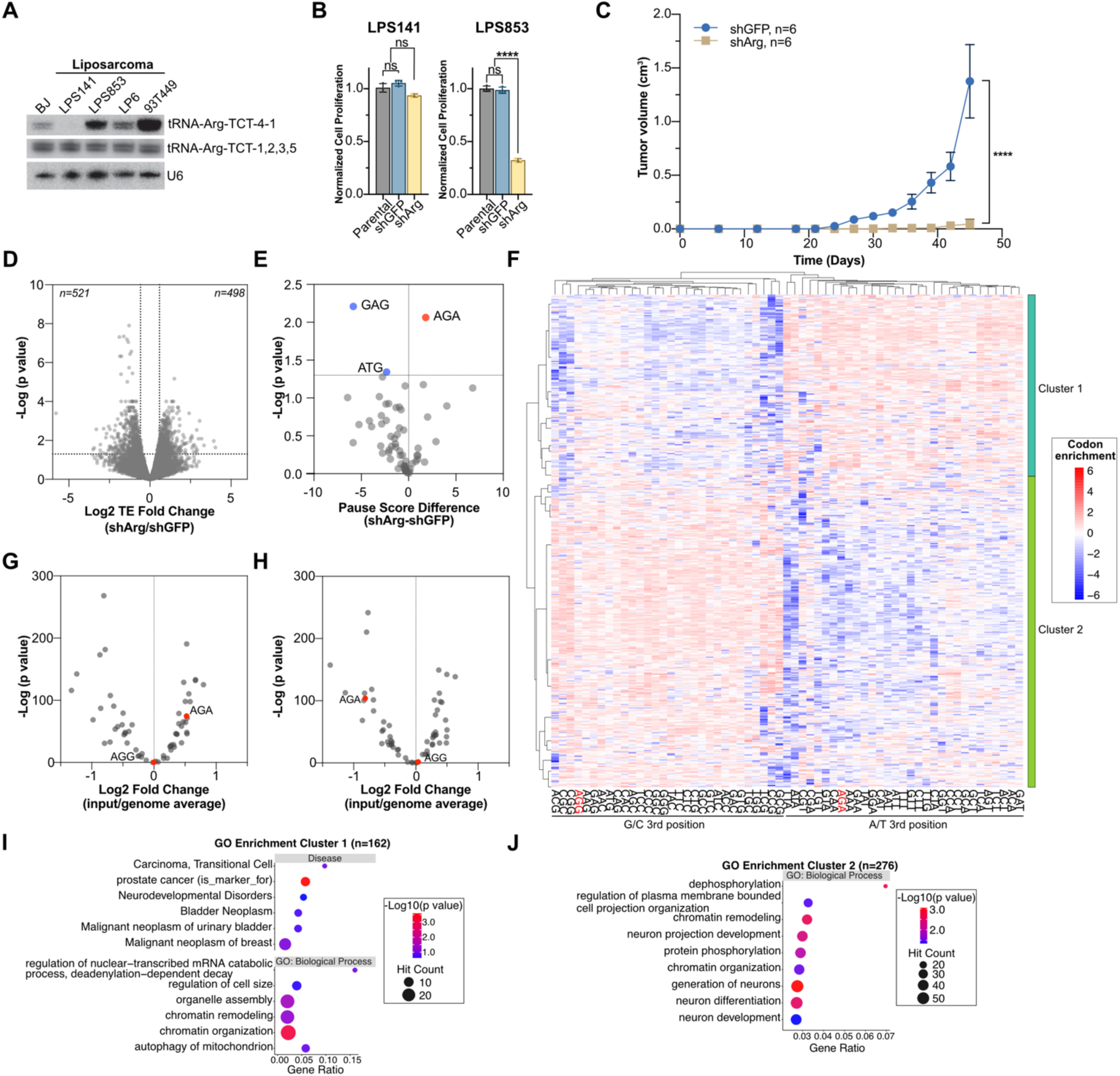
Inhibition of tRNA-Arg-TCT-4-1 in a liposarcoma model leads to altered mRNA translation and ribosome stalling. **A)** Northern blot showing relative tRNA-Arg-TCT-4-1 levels in a panel of liposarcoma cell lines. BJ: Human fibroblast. **B)** Cell proliferation of LPS141 and LPS853 cells after tRNA-Arg-TCT-4-1 inhibition. 4 days after plating. n = 3. Error bars: Mean ± S.D. P values from One-way ANOVA with posthoc Bonferroni correction. ns: not significant; ****p < 0.0001. **C)** *In vivo* tumor formation of LPS853 cells (n = 6; error bars denote mean ± SEM). ^∗∗∗∗^p < 0.0001. Two-way analysis of variance (ANOVA) and Bonferroni correction. **D)** Changes in translation efficiencies between shArg expressing cells and shGFP control cells; n = 2. **E)** Analysis of ribosome pausing between shArg and shGFP cells. P-value from Fisher’s Exact Test with Benjamini–Hochberg FDR correction. **F)** Codon enrichment analyses of genes with downregulated TE (n=521). Codon enrichment shows the fold change of each codon (61) compared to the genome-wide average. Hierarchical agglomerative clustering, with Euclidean distance and complete linkage. Codon enrichment (gene input/global genome average) of genes in cluster 1 (**G**) or cluster 2 (**H**). P-value from Fisher’s Exact Test with Benjamini–Hochberg FDR correction. Gene Ontology enrichment analysis using genes in cluster 1 (**I**) or cluster 2 (**J**) (p<0.05, TE FC≤1.5).

Subsequently, we performed SILAC proteomics, and the results show that inhibiting tRNA-Arg-TCT-4-1 leads to a broad dysregulation of the proteome in shArg-treated cells compared to the shGFP control, with most changes decreasing protein output with 576 genes significantly downregulated (<1.2-fold; p<0.05) vs 250 genes significantly upregulated (>1.2-fold; p<0.05) (**Fig. S3C, Table S3**). We then performed codon-enrichment analyses for each gene as described earlier. This resulted in the separation of the gene list into gene clusters as well as clustering by the third nucleotide (G/C or A/T) (**Fig. S3D**). Gene cluster 1 is enriched in AGA codons (**Fig. S3E**) while cluster 2 is depleted (**Fig. S3F**). Gene ontology analyses showed that the downregulated proteins are involved in mRNA processing/splicing, DNA replication/damage response, tRNA metabolism, and intracellular transport (**Fig. S3G**). Overall, these data suggest that inhibiting tRNA-Arg-TCT-4-1 in cancers outside the CNS also reduces tumor formation *in vivo*, likely due to defects in translation and ribosome stalling at AGA codons, which supports an on-target mechanism of action.

### Acute inhibition of tRNA-Arg-TCT-4-1 with antisense oligonucleotides induces cancer cell death and activates stress responses

Although our results identified tRNA-Arg-TCT-4-1 as a promising anticancer target across two cancer types, the use of shRNAs in clinical settings remains a substantial challenge. Therefore, we next opted for a different strategy to inhibit tRNA-Arg-TCT-4-1 using antisense oligonucleotides (ASOs). Mechanistically, depending on the ASO design and the nature of its interaction with the RNA substrate, it can lead to target degradation in the nucleus, mediated by RNase H, or create a steric hindrance that blocks accessibility to the target. We designed a 16-mer gapmer featuring locked nucleic acid modifications (3nt + 3nt) flanking an unmodified stretch of DNA (10nt), expecting that this could lead to tRNA degradation during its transcription and processing in the nucleus. For this, we took advantage of the small sequence differences in the body of tRNA-Arg-TCT-4-1 that would allow for the discrimination of isodecoders and specifically inhibit tRNA-Arg-TCT-4-1 (**Fig. 4A**). Next, we designed two sets of controls based on the sequence of the targeting ASO-1, a scramble sequence that contains a random sequence with the exact nucleotide content percentages and a mismatched control that contains six mismatches. As with our shRNA findings, we did not observe a decrease in tRNA-Arg-TCT-4-1 levels in cells transfected with the targeting ASO-1 (**Fig. S4A**). Nevertheless, our translation reporter system, based on the fusion mCherry-Hmga2 construct, showed a significant decrease in the translation of mCherry-Hmga2 compared to the scramble and mismatched controls (**Fig. 4B**), confirming that ASO-1 specifically inhibits tRNA function and the translation of mRNAs enriched in AGA codons. Since the genomic locus of tRNA-Arg-TCT-4-1 resides within an intronic sequence of the protein-coding gene AIM2 (**Fig. S4B**), we aimed to rule out the possibility of ASO-1 affecting AIM2 expression. To do this, we used qPCR to monitor AIM2 mRNA levels after ASO-1 treatment, compared with untreated, scrambled, and mismatched controls. The data indicate no significant changes in AIM2 levels, not only in ASO-treated cells (**Fig. S4C**) but also in shArg stable cells (**Fig. S4D**), illustrating the specificity of tRNA-Arg-TCT-4-1 targeting. After noticing translation defects in ASO-1-treated cells, we investigated whether this would affect cell proliferation. To explore this, we performed a dose titration experiment, which revealed a significant decrease in cell growth in a dose-dependent manner (**Fig. 4C**) and determined that in LPS853 cells, the IC_50_ is 16.30 ± 0.92 nM (**Fig. 4D**). Importantly, only the targeting ASO-1 (20nM) caused a significant decrease in cell proliferation compared to untreated, scramble, and mismatched controls (**Fig. 4E**). This also applies to the other two liposarcoma cell lines, LP6 and 93T449, which express tRNA-Arg-TCT-4-1 (**Fig. 3A, S4E**). To demonstrate the ASO-1 on-target specificity for the observed cell growth phenotypes, we next performed tRNA-Arg-TCT-4-1 rescue experiments. We found that modest overexpression of tRNA-Arg-TCT-4-1 (**Fig. 4F**) partially alleviates the decreased cell proliferation caused by ASO-1, further supporting an on-target mechanism (**Fig. 4G**). Because we observed a rapid decrease in cell numbers following ASO-1 treatment, we investigated whether this was due to cell death. We performed Annexin V assays to quantify the percentage of apoptotic cells post-treatment. Our results indicated that the inhibition of tRNA-Arg-TCT-4-1 leads to selective cell death in cancer cells (**Fig. 4H**) and that this response is positively correlated with isodecoder 4 expression (**Fig. 4I**). Notably, neither the non-transformed cells (BJ) nor the cancer cells with undetectable levels of the tRNA isodecoder (LPS141) were affected. Because we observed defects in translation efficiency with Ribo-Seq (**Fig. 3D**), we subsequently performed SILAC proteomics on LPS853 cells treated with ASO-1 versus ASO-Ctrl (scramble) to determine whether protein levels changed (**Table S4**). The results indicate that acute inhibition of tRNA-Arg-TCT-4-1 with ASO-1 leads to widespread proteome-wide dysregulation compared with the respective controls (**Fig. S4F**). To gain a deeper understanding, we analyzed the differentially expressed proteins. The downregulated protein list is enriched for pathways related to small-molecule, fatty acid, carbohydrate, and organic acid metabolism (**Fig. S4G**). We next reasoned that inhibition of tRNA-Arg-TCT-4-1 would cause translation defects and trigger stress responses. Consistent with this, we found that the proteins upregulated in ASO-1-treated cells are predominantly involved in processes such as translation, ribosome biogenesis, and organization of organelles and the cytoskeleton. These proteins participate in pathways including translation elongation, nonsense-mediated decay (NMD), and stress response activation due to amino acid deficiency, such as GCN2 (**Fig. S4G**). Overall, our data suggest that an ASO-mediated inhibition strategy is a viable approach to inhibiting tRNA-Arg-TCT-4-1, resulting in selectively reduced translation efficiency and cell growth defects that specifically lead to apoptosis in cancer cells expressing this tRNA isodecoder.

**Figure 4.**
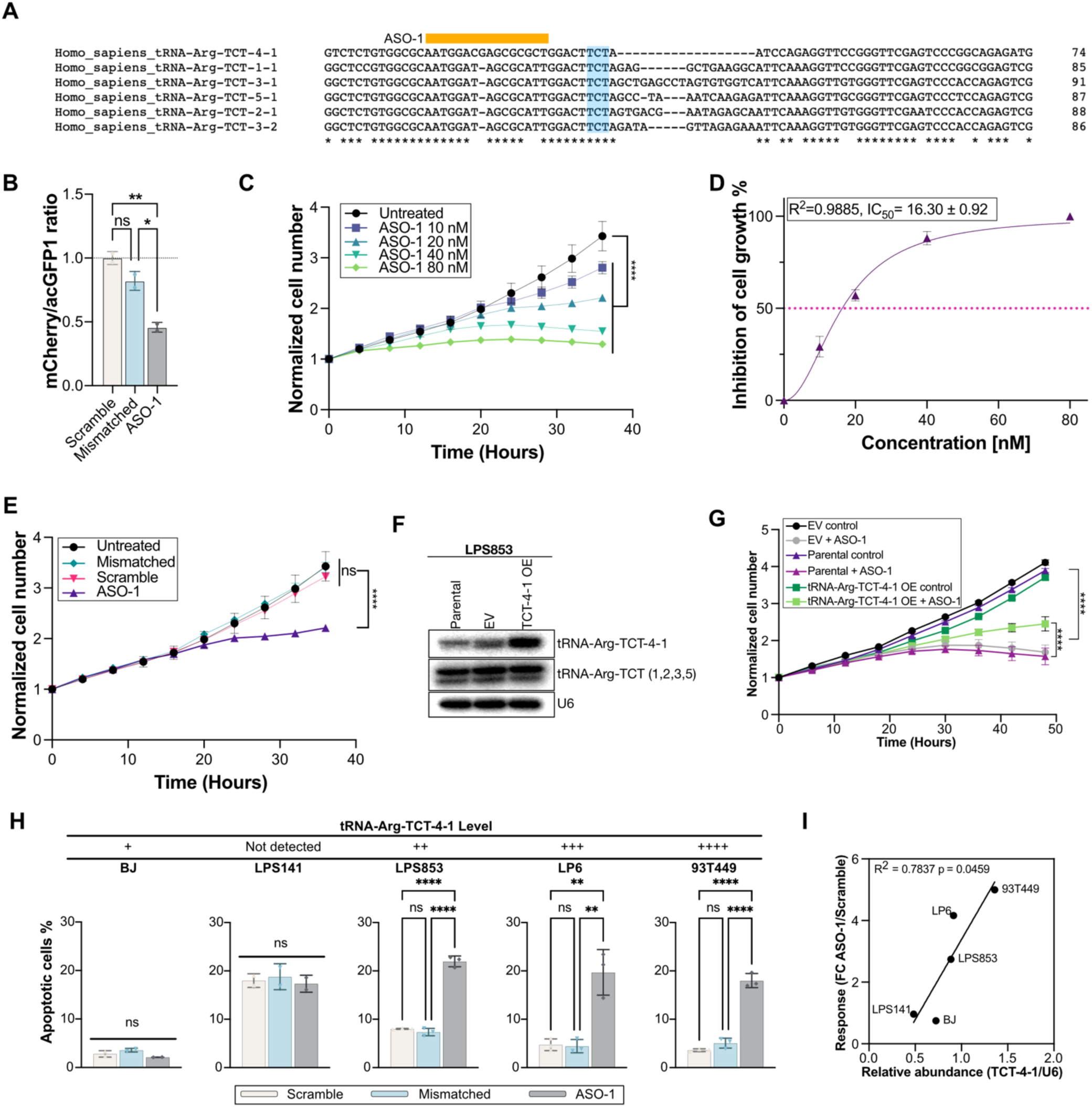
Antisense oligonucleotide-mediated tRNA-Arg-TCT-4-1 inhibition causes cell death in liposarcoma cells. **A)** Alignment of human tRNA-Arg-TCT isodecoders. Colors denote the regions targeted by ASO. Bue: anticodon region. **B)** mCherry/acGFP1 ratios measured by flow cytometry using mCherry-Hmga2-WT (5 of 12 AGA codons). Data was collected 48h post transfection. Data are mean ± SD. n = 2. p values from two-way ANOVA with Bonferroni correction. ^∗^p < 0.05 and ^∗∗^p < 0.01. ns: not significant. **C)** Normalized cell proliferation upon different concentrations of targeting ASO (ASO-1). **D)** ASO-1 dose-response and determination of IC_50_. n = 6. Error bars: Mean ± S.D. Non-linear fit with normalized response, variable slope. **E)** Normalized cell proliferation upon different ASO treatments: ASO-1, mismatched and scramble controls. 20 nM. Data are mean ± SD. n = 6. p values from two-way ANOVA with Bonferroni correction. ****p < 0.0001. ns: not significant. **F**) Northern blot showing tRNA-Arg-TCT-4-1 levels in stable cell lines. U6 is used as a loading control. **G)** Normalized cell proliferation upon ASO-1 treatment (10 nM) in parental, empty vector (EV), or tRNA-Arg-TCT-4-1 overexpressing cells. Data are mean ± SD. n = 6. p values from two-way ANOVA with Bonferroni correction. ****p < 0.0001. ns: not significant. **H)** Annexin V apoptosis assay across different cell lines expressing different tRNA-Arg-TCT-4-1 levels treated with ASO-1, scramble and mismatched controls. 20nM, 48h after transfection. n = 2 or 3. Error bars: Mean ± S.D. P values from One-way ANOVA with posthoc Bonferroni correction. ns: not significant; **p < 0.01; ****p < 0.0001. **I)** Pearson correlation analysis between normalized tRNA-Arg-TCT-4-1 abundance vs. apoptosis (response).

### Acute inhibition of tRNA-Arg-TCT-4-1 causes tumor regression *in vivo*

Encouraged by the positive results of our cellular-based assays, we next sought to investigate the effects of the acute inhibition of tRNA-Arg-TCT-4-1 on a pre-established tumor. To this end, we first tested ASO-1 activity in an LPS853 xenograft model using a single-dose intratumoral delivery of lipid nanoparticle-encapsulated ASO-1 (4mg/Kg). This resulted in a marked decrease in tumor growth compared with ASO-Ctrl (scramble) within 7 days after the injection (**Fig. 5A**). This result prompted us to pursue a larger experiment, this time on a patient-derived xenograft (PDX) model of sarcoma (SXFS627; Charles River Germany GmbH) that has high expression levels of METTL1/WDR4 (**Fig. S5A**) and detectable levels of isodecoder 4 (**Fig. S5B**). Intratumor delivery of naked ASO-1 caused tumor regression after a single 4mg/kg dose, whereas ASO-Ctrl (scramble) or vehicle controls led to rapid tumor growth, reaching humane end-point by day 16 (**Fig. 5B**). Animals treated with ASO-1 received an extra dose on day 21 and were observed for tumor growth over a period of up to 56 days. Only one animal developed a tumor on day 35, whereas the others showed no signs of tumor growth by the end of the experiment (**Fig. 5C**). Notably, no apparent toxicity or weight loss was observed (**Fig. 5D**). Taken together, the data show that inhibiting tRNA-Arg-TCT-4-1 in established tumors induces tumor regression and extends survival, with no apparent toxicity.

**Figure 5.**
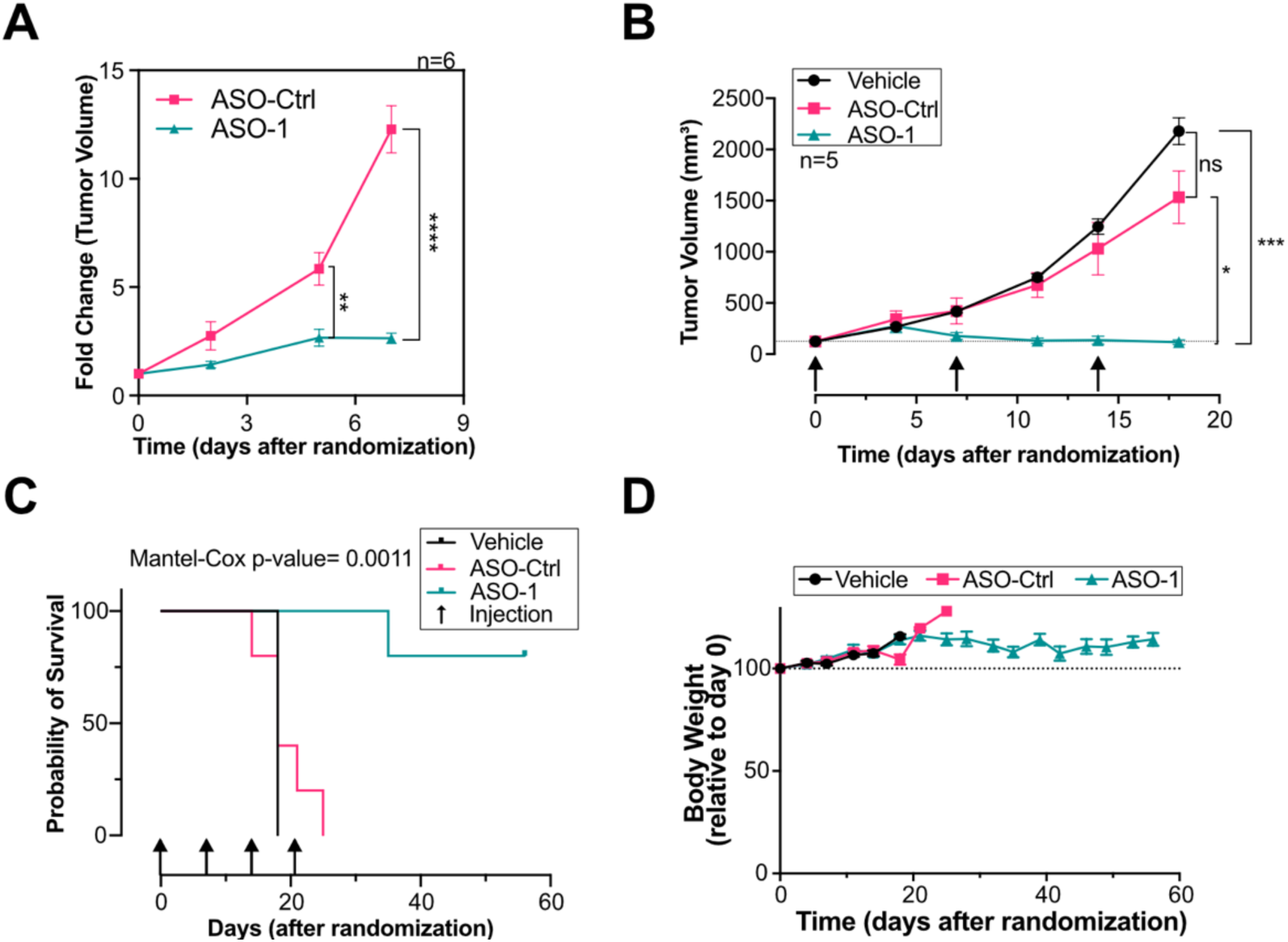
Acute inhibition of tRNA-Arg-TCT-4-1 causes tumor regression and extends survival. **A)** Effect on LPS853 tumor growth upon a single ASO-1 treatment (4mg/Kg). Average initial volume of 100 mm^3^. Data are mean ± SD. n = 6. p values from two-way ANOVA with Bonferroni correction. ****p < 0.0001; **p < 0.01. **B)** Effect on SXFS627 (PDX) tumor growth upon ASO-1 treatment (4mg/Kg). Average initial volume of 100 mm^3^. Data are mean ± SD. n = 5. P-values from two-way ANOVA with Bonferroni correction. ****p < 0.0001; **p < 0.01. Arrows indicate injection days. **C)** Kaplan-Meier survival curve of animals from **(B)**. P-value from Mantel-Cox test. **D)** Body weight changes in animals treated with vehicle (PBS), ASO-1 or Ctrl (scramble). n = 5.

## Discussion

This study underscores the critical role of tRNA-Arg-TCT-4-1 in cancer biology, highlighting its potential as a novel therapeutic target. Our findings demonstrate that tRNA-Arg-TCT-4-1 is consistently overexpressed in a variety of cancer types, with its highest levels observed in aggressive forms of cancer such as breast-invasive adenocarcinoma and glioblastoma. This overexpression is closely associated with poor patient outcomes, suggesting that tRNA-Arg-TCT-4-1 plays a significant role in oncogenesis. Mechanistically, we have previously shown that the oncogenic potential of tRNA-Arg-TCT-4-1 is mediated through its ability to selectively enhance the translation of proteins enriched with AGA codons(*10*). These genes are involved in critical pathways such as cell cycle regulation, metabolic adaptation, and stress responses, processes that are essential for tumor growth and survival. The upregulation of tRNA-Arg-TCT-4-1 is driven by both epigenetic and post-transcriptional mechanisms, as evidenced by the hypomethylation of its encoding DNA region and METTL1-mediated m^7^G stabilization (*10*, *39*). These findings align with prior studies implicating METTL1 in reshaping the tRNA pool to promote oncogenic translation reprogramming.

To our knowledge, this is the first time that inhibiting a tRNA isodecoder has been demonstrated to have potential therapeutic applications. Other studies have focused on studying tRNA roles by inhibiting a whole tRNA family (*13*). However, an important observation in this study is the apparent inability of other tRNA-Arg-TCT isodecoders to compensate for the deficiency of tRNA-Arg-TCT-4-1, as evidenced by the strong molecular and cellular phenotypes upon tRNA inhibition in different human cancer cells and PDX models. Focusing on tRNA isodecoder-specific functions could, therefore, unveil new dimensions of translational control in cancer and other diseases, offering opportunities for highly targeted therapeutic interventions.

Importantly, this study demonstrates that targeted inhibition of tRNA-Arg-TCT-4-1, either through shRNA or antisense oligonucleotides (ASOs), effectively suppresses tumor formation and causes tumor regression *in vivo*. These approaches lead to significant translational defects, including reduced efficiency in synthesizing proteins translated from AGA-enriched mRNAs and increased ribosome stalling at AGA codons. The resultant proteomic dysregulation particularly affects pathways related to RNA metabolism, cellular metabolism, and stress responses, emphasizing the central role of tRNA-Arg-TCT-4-1 in maintaining the malignant phenotype. The specificity of tRNA-Arg-TCT-4-1 targeting is a significant advantage. The selective overexpression of this isodecoder in cancer cells, coupled with its minimal expression in normal tissues, predominantly restricted to the central nervous system(*38*, *43*), supports the notion that therapeutic strategies targeting this tRNA would have an exploitable therapeutic window. Furthermore, the protective role of the blood-brain barrier for its physiological expression sites could help mitigate concerns about potential adverse effects on normal tissues, and future work is needed to understand the safety profile of the inhibition strategy.

We hypothesize that ASO-mediated inhibition of tRNA-Arg-TCT-4-1 is driven by steric hindrance (sponging). Unlike canonical RNA interference (RNAi) approaches, which often rely on degradation of the RNA target (when there is 100% base complementarity)(*44*), ASOs designed for steric hindrance physically obstruct the functional domains of the target molecule. This prevents the interaction of the tRNA with its associated machinery, potentially including aminoacyl-tRNA synthetases, ribosomes, and elongation factors. This blockage leads to stalled ribosomes, often considered toxic for cells(*45*, *46*). Ribosome stalling activates the ribotoxic stress response (RSR)(*47*) and downstream integrated stress response pathways, including the responses to amino acid deficiency and activation of the NMD pathway, as cells attempt to mitigate the burden of misfolded and incomplete proteins. These stress responses ultimately lead to apoptotic cell death, as evidenced by increased Annexin-V staining in ASO-1-treated cancer cells. Importantly, these effects were specific to cells expressing detectable levels of tRNA-Arg-TCT-4-1, underscoring the precision of this therapeutic approach.

While the findings of this study are highly promising, several questions remain to be addressed. The structural and functional features that confer oncogenic properties to tRNA-Arg-TCT-4-1 warrant further investigation. Additionally, optimizing ASO delivery systems to improve tumor targeting and minimize immune responses to the ASO will be critical for successful clinical translation. Exploring the interactions between tRNA-targeted therapies and existing cancer treatments, particularly those targeting the translation machinery or epigenetic regulators, could reveal synergistic effects and broaden therapeutic options. One aspect that could be particularly exciting is the development of small-molecule inhibitors that bind tRNA-Arg-TCT-4-1. We have shown that target depletion is not necessary for tumor suppression, and that steric hindrance may be sufficient to exert a therapeutic effect. Due to an initial hypothesis that tRNA isodecoders are simply an evolutionary relic(*48*) and the fact that tRNAs have a conserved shape(*49*), the molecular details of their structures, folding, and dynamics have been significantly understudied. This lack of structural information is currently one of the challenges impeding the development of chemical inhibitors. While there are no published preclinical data to support the use of tRNAs as treatment targets for diseases to date, there is some precedent from an accidental discovery(*50*). The small molecule Velcrin promotes apoptosis by forming a heterotetramer with PDE3A, velcrin, and SLFN12, leading to targeted degradation of tRNA-Leu-TAA in Velcrin-sensitive cells(*51*). Targeting structured RNAs with small molecules has recently shown significant advantages (*52*). Often, these studies use tRNAs as controls, inadvertently identifying hits that bind to tRNAs, thereby demonstrating the potential to identify specific tRNA binders. We anticipate that with the increased attention the field of tRNA biology has received in recent years, new therapies targeting tRNAs will emerge in the future.

In conclusion, this study establishes tRNA-Arg-TCT-4-1 as a pivotal driver of oncogenesis and a promising therapeutic target. By elucidating its role in cancer progression and demonstrating the feasibility of its inhibition, these findings pave the way for innovative tRNA-focused cancer therapies. By specifically targeting oncogenic isodecoders like tRNA-Arg-TCT-4-1, it is possible to induce translational arrest and activate stress responses in cancer cells while sparing normal tissues. This strategy not only broadens the scope of RNA-targeted therapies but also challenges long-held assumptions about the functional redundancy of tRNA isoforms. The insights from this research have the potential to significantly advance precision medicine and improve patient outcomes across a diverse range of cancer types.

## Materials and Methods

### Cell Lines

Human glioblastoma LNZ308 cells (male) (CRL11543) were purchased from ATCC. LP6 (sex unspecified)(*53*) was a gift from Eric Snyder, LPS141 (sex unspecified) (*53*) and LPS853 (sex unspecified)(*54*) were gifts from Jonathan Fletcher, and 93T449 (female)(*55*) was a gift from Florence Pedeutour. LNZ308 were cultured in DMEM supplemented with 10% FBS and 1X penicillin/streptomycin. LPS141 and 93T449 cells were cultured in RPMI 1640 medium supplemented with 15% FBS and 1X penicillin/streptomycin. LPS853 cells were cultured in IMDM medium supplemented with 15% FBS, 1X GlutaMax and 1X penicillin/streptomycin. LP6 cells were cultured in DMEM/F12 medium supplemented with 10% FBS, 1% GlutaMax and 1X penicillin/streptomycin. All cell lines were cultured in the presence of 5% CO_2_ at 37°C. Human cell lines employed are not listed in the cross-contaminated or misidentified cell lines database curated by the International Cell Line Authentication Committee (ICLAC). All cell lines tested negative for mycoplasma contamination.

### Animal subjects

4-6 weeks old female NU/J (Nude) or NOD scid gamma (NSG) immunodeficient mice (Jackson Laboratory) were used for subcutaneous injections.

### Plasmid Construction

Individual tRNAs were overexpressed as previously reported(*10*). To generate a short hairpin targeting tRNA-Arg-TCT-4-1 (shArg), we used a pLKO.1-Puro backbone. We first annealed our shRNA oligos and inserted them into a linearized pLKO.1-Puro backbone using EcoRI and AgeI restriction sites. Clones were screened and submitted for Sanger sequencing for correct insertion validation. The sequences used for shRNA cloning are listed in **Table S5.**

### Virus production and generation of stable knockdown and overexpression cells

Generation of stable knockdown and overexpression cells *via* virus transduction was performed as described previously(*56*). In brief, shRNA containing pLKO.1 vector was co-transfected with pLP1, pLP2, and VSVG into 293T cells. Viruses were collected at 48 h and 72 h after transfection and then used to infect cells; 48 h after infection, puromycin (2.5 ug/mL) was added to the culture medium to select the infected cells. LNZ308, and LPS853 cells infected with shARG, or shGFP were maintained in medium with puromycin (2.5 ug/ml). The medium was refreshed on the following day and the transduced cells were cultured further.

### ASO transfection and cell proliferation assays

LPS853 cells expressing fluorescent proteins mCherry (nuclear) and acGFP were maintained at 37°C and 5% CO_2_ in complete media containing Iscove’s Modified Dulbecco’s Medium (IMDM), supplemented with 15% Fetal Bovine Serum (FBS), 1X GlutaMax, and 1X Penicillin/streptomycin. To test the effect of ASO treatment on cell proliferation, LPS853 cells were transfected with a targeting ASO-1, along with non-targeting controls ASO mismatch (MM) and scramble (SC) in a 96-well plate. All ASOs were purchased from IDT. Each well was seeded with cells in 50 µL of complete media 24 hours prior to the transfection and incubated at 37°C and 5% CO_2_. The next day, for the ASO treatment, 10 nM, 20 nM, 40 nM, and 80 nM of each ASO were transfected into the cells using 0.2 µL of Lipofectamine RNAiMax reagent per well. Cell proliferation assays were conducted over 36 hours in a BioTek Cytation 5 Cell Imaging Multimode Reader using a GFP and Texas Red imaging filter with a 10X objective, maintaining low illumination intensity every 4 or 6 hours, and data was normalized both to time zero and to untreated cell controls. For rescue experiments, LPS853 was transduced with a lentivirus encoding tRNA-Arg-TCT-4-1 as described previously(*2*). The ASO sequences are listed in **Table S5.**

### Soft agar colony formation assays

Soft agar colony formation assays were performed as previously described(*10*). Briefly, fifty thousand single live MEFs cells were mixed with 0.35% top-agar (SeaPlaque, Lonza) and were plated onto 0.7% base-agar (SeaPlaque, Lonza) in six-well plates. Twenty-eight days after plating the cells into soft agar, colony numbers were counted. The plates were imaged using a EVOS FL auto plate imager (Thermo Fisher Scientific) under continuous scan. Images were stitched and the colony numbers were counted using ImageJ.

### Fluorescent reporter assay

Fluorescent reporter assays were performed as previously described(*10*). In brief, 8×10^5^ LPS853 or LNZ308 cells expressing shGFP or shArg were seeded in a 60mm plate on day 0. On day 1, cells were transfected with 2µg of mCherry-Hmga2-WT or mCherry-Hmga2-MUT using Lipofectamine 2000 (Life Technologies) and 48 hours later cells were trypsinized and resuspended in fresh medium to a density of 1×10^6^ cells/mL. For ASO testing, cells were co-transfected with 20nM ASO plus 2µg of mCherry-Hmga2-WT or mCherry-Hmga2-MUT using Lipofectamine 2000 (Life Technologies) and processed as described above. Cells were analyzed using the BDFortessa LSRII Cell Analyzer (BD Pharmigen) and acGFP1 and mCherry expression was monitored. Data were analyzed using FlowJo V10.7 (Beckton Dickinson and Co.) and presented as a ratio between mCherry and acGFP1 intensity levels.

### Quantitative RT–qPCR

Total RNA was isolated from cancer cells using Trizol. For cDNA synthesis, total RNA was reverse-transcribed with the SuperScript III cDNA Synthesis kit (Life Technologies). The levels of specific RNAs were measured using a StepOne real-time PCR machine and the Fast SybrGreen PCR mastermix (ThermoFisher, 4385612) according to the manufacturer’s instructions. All samples, including the template controls, were assayed in triplicate. The relative number of target transcripts was normalized to β-Actin. The relative quantification of target gene expression was performed with the comparative cycle threshold (*C*_T_) method.

### Quantitative tRNA RT–qPCR

To remove RNA modifications that block primer extension upon reverse transcription, total RNA isolated from the four LPS cell lines (LPS141, LP6, LPS853, and 93T499) and total RNA from PDX tumor samples were treated in the presence of AlkB and mutant AlkB D135S in 1X demethylation buffer (final 300mM KCl, 2mM MgCl_2_, 50µM ammonium iron (II) sulfate hexahydrate, 300µM 2-ketoglutarate, 2mM L-asorbic acid, 50µg/mL BSA, 50mM MES pH 5, and RNasIn (Promega) ribonuclease inhibitor) at room temperature for 2 hours. Wild-type AlkB was used at 5x the amount of total RNA and mutant D135S AlkB was used at 10x the amount of final RNA (up to 10 µg). Reactions were quenched with 1µL 0.5M EDTA pH8 and 2µL DNaseI (New England Biolabs) added and incubated at 37°C for 15 minutes. RNA was cleaned up and isolated using the RNA clean and concentrator kit (Zymo Research) and in-column DNase treatment also performed. To degrade lingering aminoacyl bonds, the cleaned-up RNA was incubated in 1M Tris pH9 for one hour at 37°C followed by cleanup using the RNA clean and concentrator kit. RNA was polyadenylated using 5 units *E. coli* Poly(A) Polymerase (New England Biolabs) in final concentration 1mM ATP and 1X *E. coli* Poly(A) Polymerase reaction buffer and incubated at 37° C for one hour followed by the RNA clean and concentrator kit. Total RNA was used as template in the reverse transcription reaction to create cDNA. RNA was incubated with 5 µM (final) oligo dT20 and 1 µM (final) dNTPs for 65°C for 5 minutes and allowed to cool to 4°C. Added to this reaction was a mixture of the following: 10X RT buffer, 25mM MgCl_2_, 0.1M DTT, 1 µL RNaseOut, and 1uL SuperScript III (ThermoFisher). To yield cDNA, this reaction was incubated at 50°C for 50 minutes followed by enzyme inactivation at 85°C for 5 minutes. 1 µL of RNaseH was subsequently added and the mixture incubated at 37°C for 20 minutes to degrade RNA template. cDNA stocks were diluted with water to 5ng/ µL.

To quantify levels of tRNA-Arg-TCT-4-1, 10ng of cDNA was mixed with final concentration 1X Luna Universal qPCR Master Mix (NEB), 0.25µM Forward Primer, 0.25µM Reverse Primer, and 0.2µM FAM-Taqman probe (**Table S5**). Samples were run in triplicate at 95°C for 3 minutes followed by 45 cycles of denaturation at 95°C for 10 seconds and extension at 60°C for 30 seconds. To extrapolate the number of tRNA Arg TCT 4-1 molecules in each tumor sample and liposarcoma cell line, a standard curve of tRNA Arg TCT 4-1 was created. To create the standard curve, cDNA from *in vitro* tRNA Arg TCT 4-1 was used as the template in qPCR reactions under the same conditions described above. The calculated molecules of tRNA per ng of input template plasmid used in each qPCR reaction were plotted versus the average Cq value from the qPCR reaction to create a standard curve. The slope and y-intercept of the standard curve were then used to calculate the number of tRNA molecules in each tumor and liposarcoma cell line sample, based on their average Cq values, which fell within the standard curve’s range.

### tRNA sequencing

tRNA sequencing was performed as previously described(*56*, *57*). First, isolated small RNAs were treated with recombinant wild-type and D135S AlkB proteins to remove the dominant methylations on RNAs. 10 µg small RNAs were treated with 80 pmol wt AlkB and 160 pmol D135S AlkB mutant for 2 hours in a 100 µl demethylation reaction [300 mM KCl, 2 mM MgCl_2_, 50 mM of (NH_4_)_2_Fe(SO_4_)2·6H_2_O, 300 mM 2-ketoglutarate (2-KG), 2 mM L-ascorbic acid, 50 mg/mL BSA, 50 mM MES buffer (pH 5.0)] at room temperature. After incubation, the reaction was quenched with a final concentration of 5 mM EDTA and the RNAs were purified by phenol–chloroform extraction followed by ethanol precipitation. Alkb-treated RNAs (2.5 ug) were then treated with 0.1M NaBH_4_ for 30 min on ice at dark in the presence of 1 mM free m^7^GTP as methylation carrier. RNAs were precipitated with sodium acetate (300mM final concentration, pH5.2) and 2.5 volumes of cold ethanol at −20C overnight. The NaBH_4_-treated RNAs were subsequently treated with aniline-acetate solution (H_2_O: glacial acetate acid:aniline, 7:3:1) at room temperature at dark for 2 hours to induce the site-specific cleavage. The RNA samples were purified by ethanol precipitation and used for cDNA library construction using NEBNext Small RNA Library Prep Set (New England Biolabs) followed by sequencing with Illumina Nextseq 500.

### TCGA data analysis

TCGA analyses were performed as previously described(*10*) RNA-Seq expression data and small RNA-Seq data for 33 TCGA tumor types were downloaded from the Genomic Data Commons Data Portal (GDC) of TCGA (http://cancergenome.nih.gov/). The gene expression matrix was then constructed by merging the TPM (Transcripts Per Million) values of all RNA-seq samples. The tumor types with normal tissues were used to draw the gene expression boxplots and the statistic differences are then calculated using Wilcoxon rank-sum test (with asterisks indicating statistical significance). The expression correlations between METTL1 and WDR4 among TCGA tumors were conducted by using Pearson correlation coefficient. The tRNA expressions from TCGA small RNA-seq data were analyzed using ARM-Seq data analysis pipeline(*58*). We used data generated by the Clinical Proteomic Tumor Analysis Consortium (NCI/NIH) to analyze METTL1 protein levels versus RNA transcripts.

### Ribosome footprinting (Ribo-seq)

Cancer cells were grown to 80%–90% confluence in DMEM supplemented with 10% FBS in 15-cm dishes. Ribosome footprinting was performed according to TruSeq® Ribo Profile system (Illumina) with modifications. Briefly, cells were treated with 0.1 mg/mL cycloheximide (CHX) for 1 minute to inhibit translation elongation and washed with ice-cold PBS containing 0.1 mg/mL CHX. Next, cells were lysed in 800 μl 1X Mammalian Polysome Buffer (Illumina) supplemented with 1% Triton X-100, 1 mM DTT, 10 units DNase I, 0.1 mg/mL CHX, and 0.1% NP-40. Lysates were cleared by centrifugation at 12,000 g for 10 min at 4°C, and supernatants were flash-frozen in liquid nitrogen and stored at −80°C until processing. RNA concentration of the lysates was measured according to their absorbance at A_260_ and an equivalent of 12xA_260_/ml was treated with 5 U/A_260_ TruSeq Ribo Profile RNase Nuclease (Illumina) for 45 minutes at room temperature. RNase activity was inhibited by adding 15 μl SUPERase to the mixture. Ribosome protected fragments (RPFs) were isolated using MicroSpin S-400 columns (GE Healthcare). RPF RNA samples (5μg) were subjected to ribosomal RNA depletion using RiboMinus Eukaryote Kit v2 (Thermo Fisher Scientific). Ribo-depleted RNA samples were separated on 15% polyacrylamide/TBE/Urea gels (Thermo Fisher Scientific) and the RNA fragments corresponding to ∼25-35 nt were excised. RNA was gel-extracted and precipitated overnight at 4°C using 0.5 M ammonium acetate. In parallel, total RNA input samples were isolated, and fragmented at 94°C for 25 minutes. Input total RNA and RPFs were subjected to end repair by TruSeq Ribo Profile PNK (Illumina), cleaned using RNA Clean & Concentrator-5 kit (ZYMO Research) and ligated to 2.5 μM Universal miRNA Cloning Linker (NEB) by using 100 units T4 RNA Ligase 2, truncated KQ (NEB) for 3 hours at 22°C. After ligation, RNA samples from both total RNA and RPFs were reverse transcribed using SuperScript III Reverse Transcriptase (Thermo Fisher Scientific) and 0.25 μM RT primer (IDT). cDNA samples were then gel-extracted on 10% polyacrylamide/TBE/Urea gels (Thermo Fisher Scientific) and circularized by CircLigase ssDNA Ligase (Lucigen) for 2 hours at 60°C. cDNA libraries for total RNA and RPFs were amplified for 9 and 12 PCR cycles, respectively, using Phusion High-Fidelity PCR Master Mix (NEB), Illumina index primers and 10 μM forward primer (IDT). Amplified libraries were cleaned using AMPure XP Beads (Beckman Coulter), followed by gel-extraction on 8% native TBE gels (Thermo Fisher Scientific). Libraries were sequenced with Illumina NextSeq 500.

### Ribo-Seq data analysis

The sequences of input and Ribo-seq samples were firstly processed to get the clean reads by trimming the adapters and filtering the low-quality sequences. Then, for Ribo-seq input data, the clean reads were aligned to reference genome sequences using STAR (*59*). The resulted BAM mapping files were used as inputs of HTSeq (*60*) to calculate the read counts for each gene from GENCODE gene mode. For the cleaned Ribo-seq data, the clean ribosome-protected fragments (RPFs) were firstly collapsed into FASTA format by fq2collapedFa. RiboToolkit (https://bioinformatics.caf.ac.cn/RiboToolkit_demo) was used to perform codon occupancy analysis and translation efficiency analysis by uploading the collapsed RPF tags and gene read counts from input samples(*61*). In brief, rRNA and tRNA sequences were filtered from RPF containing files by alignment to rRNA sequences (Ensembl non-coding, release 91) (*62*) and tRNA sequences from GtRNAdb databases(*63*). The resulting ribosome-protected fragments (RPFs) were aligned to the mouse reference genome (mm10) using STAR [10.1093/bioinformatics/bts635] and only unique mapped reads were kept. The genome unique mapping reads were then mapped to transcript sequences using Bowtie with a maximum of one mismatch allowed(*64*) and all the transcript mappings were kept. CONCUR tool (https://github.com/susbo/concur) was used for calculating codon usage based on the reference genome mapping based on STAR(*59*, *65*). The translation efficiency (TE) was calculated by dividing RPF abundance on CDS by its mRNA abundance of input sample. A threshold of two-fold change and FDR <0.05 was used to define the differential translation genes (*66*). PausePred (https://pausepred.ucc.ie/), was used to infer ribosome pauses from Ribo-seq data. Peaks of ribosome footprint density are scored based on their magnitude relative to the background density within the surrounding area (*67*).

### Proteomic analysis by stable isotope labeling using amino acids in cell culture (SILAC)

LNZ308 human glioblastoma cells or LPS853 liposarcoma cells were grown in media supplemented with isotopic-labeled ^13^C6^15^N2 l-lysine and ^13^C6^15^N4 l-arginine (heavy) or normal amino acids (light) for 15 to 21 days until a labeling efficiency >95% was achieved following the instructions of the SILAC Protein Quantitation Kit (Trypsin) – DMEM (A33972, Thermo Scientific). Cells were lysed in 1x passive cell lysis buffer (Promega) supplemented with cOmplete protease inhibitor (11873580001, Roche). Protein lysates were cleared by centrifugation at 14,000xg for 5 minutes at 4°C. Equal amounts of heavy and light protein (1:1) amounts were mixed with 2x reducing sample buffer and 100 µg of clarified sample was separated by SDS-PAGE (4-20%). Three technical replicates were performed for each sample. Gels were stained using Novex Colloidal blue staining (Invitrogen) and each lane was cut into 12 slices of equal size. Excised gel bands were cut into approximately 1 mm^3^ pieces. The samples were reduced with 1 mM DTT for 30 minutes at 60°C and then alkylated with 5mM iodoacetamide for 15 minutes in the dark at room temperature. Gel pieces were subjected to a modified in-gel trypsin digestion procedure(*68*). Gel pieces were washed and dehydrated with acetonitrile for 10 min. followed by removal of acetonitrile. Pieces were then completely dried in a speed-vac. Rehydration of the gel pieces was with 50 mM ammonium bicarbonate solution containing 12.5 ng/µl modified sequencing-grade trypsin (Promega, Madison, WI) at 4°C. Samples were then placed in a 37°C room overnight. Peptides were later extracted by removing the ammonium bicarbonate solution, followed by one wash with a solution containing 50% acetonitrile and 1% formic acid. The extracts were then dried in a speed-vac (∼1 hr). The samples were then stored at 4°C until analysis. On the day of analysis, the samples were reconstituted in 5 - 10 µl of HPLC solvent A (2.5% acetonitrile, 0.1% formic acid). A nano-scale reverse-phase HPLC capillary column was created by packing 2.6 µm C18 spherical silica beads into a fused silica capillary (100 µm inner diameter x ∼30 cm length) with a flame-drawn tip (*69*). After equilibrating the column, each sample was loaded via a Famos auto sampler (LC Packings, San Francisco CA) onto the column. A gradient was formed, and peptides were eluted with increasing concentrations of solvent B (97.5% acetonitrile, 0.1% formic acid). As each peptide was eluted, they were subjected to electrospray ionization and then entered into an LTQ Orbitrap Velos Pro ion-trap mass spectrometer (Thermo Fisher Scientific, San Jose, CA). Eluting peptides were detected, isolated, and fragmented to produce a tandem mass spectrum of specific fragment ions for each peptide. Peptide sequences (and hence protein identity) were determined by matching protein or translated nucleotide databases with the acquired fragmentation pattern by the software program, Sequest (ThermoFinnigan, San Jose, CA)(*70*). The differential modification of 8.0142 and 10.0083 mass units for lysine and arginine, respectively were included in the database searches to find SILAC labeled peptides. All databases include a reversed version of all the sequences and the data was filtered to between a one percent or lower peptide false discovery rate. SILAC protein ratios (H/L) were determined as the average of all peptide ratios assigned to a protein between heavy and light samples. Differential protein expression was determined using a moderated t-test, testing for the null-hypothesis being no change in H/L ratio. Multiple tests were corrected using false discovery rate (<1%).

### Analyses of codon bias

Codon usage enrichment was calculated on a per-gene basis using canonical transcript-level codon count data. Codon counts were aggregated across all transcripts for each gene of interest, excluding stop codons and non-codon metadata columns. Genome-wide background codon frequencies were computed from all canonical transcripts to serve as a reference distribution (GRCh37). For each gene, codon frequencies were compared to background frequencies, and enrichment was quantified as log2 fold change and enrichment ratios with a small pseudocount to avoid division by zero. Statistical significance was assessed for each codon using Fisher’s exact test, followed by Benjamini–Hochberg correction for multiple testing. The resulting enrichment matrix was cleaned to remove duplicates, missing values, and zero-variance rows or columns, and extreme values were capped for visualization stability. Hierarchical clustering of genes based on their codon enrichment profiles was performed, and clustered heatmaps were generated to visualize patterns of codon usage bias. Gene clusters were extracted and saved, and both pheatmap- and ggplot2-based heatmaps were produced to facilitate the interpretation of codon enrichment patterns across genes.

### Northern Blot

For Northern blotting of tRNAs or U6 snoRNA, 10µg total RNA samples were mixed with 2X TBE loading buffer (Bio-Rad) and incubated at 95°C for 5 min. The samples were then loaded into 15% TBE-UREA (Bio-Rad) gels to separate the RNAs by molecular weight. Next, the RNAs were transferred onto a positively charged nylon membrane and crosslinked with UV. For Northern blots, the membrane was blotted with radioactive labeled probes against tRNAs or U6 snRNA. Acid urea PAGE (10%) was used to evaluate aminoacylation levels in the presence of 10 mM CuSO_4_ or when the pH was above 9 (alkaline) or below 6 (acidic) pH. The probe sequences are listed in **Table S5**. Blotted membranes were then exposed to autoradiography films or phosphor-screens.

### Animal studies

4-6 week old female NU/J (Nude) immunodeficient mice (Jackson Laboratory #002019) were used for subcutaneous injections. LNZ308 or LPS853 cells (5×10^6^ cells) shGFP or shARG were transplanted into nude mice. The indicated number of cells were mixed with serum-free medium and growth factor reduced Matrigel (Corning #354230) (1:1) and injected into the right flank of nude mice. Five or six mice were used for each group. Subcutaneous tumor formation was monitored by calipers twice a week. The tumor volume was calculated using the formula 1/2(length × width^2^). The recipient mice were monitored and euthanized when the tumors reached 1 cm in diameter. At end-point, tumors were collected and weighed. Randomization and blinding were not applied. For ASO-1 experiments, LPS853 cells (5×10^6^ cells) were transplanted into nude mice as described above. Six mice were used for each group. Subcutaneous tumor formation was monitored by calipers twice a week, and once the tumors reached an average of 100 mm^³^, animals were randomized into two groups. ASO-1 or Control (scramble) was prepared using Invivofectamine (Invitrogen) following the instructions of the manufacturer, and single intratumor injections (4mg/mL) were administered once and tumor volumes were monitored for a week. The experiment was terminated when tumors reached 1,500 – 2,000 mm^³^.

The PDX model used in this study, SXFS627, was developed by Charles River Laboratories Germany GmbH, and the experiments were conducted under contract. The SXFS627 model was established from a recurrent rhabdomyosarcoma sample from a 55-year-old female patient and was selected based on METTL1 and WDR4 expression using available data from Charles River Laboratories. SXFS627 tumors were transplanted into NSG female mice. Five mice were used for each group. Subcutaneous tumor growth was monitored with calipers twice weekly, and once tumors reached an average of 100 mm³, animals were randomized into three groups so that all groups have the same average tumor volume. Animals were dosed with vehicle (PBS), ASO-1, or Control (scramble) (4 mg/mL) via intratumoral delivery. Dosing occurred once every 7 days, and tumor volumes were estimated as described above. The experiment was terminated when tumors reached 1,500 – 2,000 mm^³^.

### tRNA Fluorescent In-situ Hybridization (FISH) using BaseScope

Paraffinized breast cancer and liposarcoma (BR725 and SO2083a) human tissue microarrays were obtained from TissueArray. The ACD BaseScope protocol for preparing FFPE slides for the RNAscope 2.5 assay was carried out with minor modifications. ACDBio designed a probe specific for tRNA-Arg-TCT-4-1 (Cat. 1319851-C1). The specificity of the probe was first determined using a mouse (B6N) cerebellum. Histoclear was utilized instead of xylene for deparaffinizing the slides. For hydrogen peroxide (H_2_O_2_) incubation, the slides were incubated four times for 10 minutes each with fresh 6% H_2_O_2_ diluted in 1X PBS, rather than the RNAscope hydrogen peroxide. The Manual Target Retrieval Protocol involved boiling the slides for five minutes in Target Retrieval Reagent, transferring them to room temperature water for one minute, and boiling them again for an additional four minutes. Slides were incubated in RNAscope Protease Plus for 30 minutes. For the BaseScope v2 Assay, all steps were conducted according to the protocol, except for amplification step 7, which was left overnight instead of for two hours. After following the BaseScope protocol, the ThermoFisher Scientific DAPI protocol for fluorescence imaging was adhered to, except the slides were incubated for 15 minutes with 300 nM DAPI. A Cytation5 instrument was employed to image the entire tissue array with Texas Red and DAPI settings using a 10X zoom.

For individual tissue arrays, 60X confocal images were captured on a spinning confocal disk under the same laser power and exposure times, using a Zyla camera with NIS Elements Advanced Research Software 5.11.02. The images were processed using ImageJ. To quantify the Texas red (TR) signal of tRNA-Arg-TCT-4-1 levels in each tissue array, a modified objective scale from 0 to 4 was used based on the Basescope protocol, with each number indicating the following. 0: no TR signal or containing less than one tRNA puncta to every 20 cells, 1: cells with signal have one puncta per cell, 2: cells with signal have at least 2-3 puncta but very few clusters of puncta, 3: ≤10% of cells with puncta contain clusters, and 4: ≥10% of cells with puncta contain clusters. Data were plotted using Prism, and one-way ANOVA with Bonferroni correction analysis was performed to determine significant differences.

### Quantification and statistical analysis

Quantification and statistical analysis methods were described in individual method sections and Figure legends. Center is represented by mean and dispersion is represented by the standard deviation. The n is reported in each figure legend. Paired Student’s T tests, were used for two group comparisons. One-way ANOVA was used for multigroup comparison. Bonferroni post-hoc p-value corrections were performed for multigroup comparisons. Pearson tests were performed for correlation analysis. Randomization and blinding were not applied. The ROUT method was used to identify outliers (Q = 1%). Data were considered significant if p-values < 0.05.

## Acknowledgments

E.A.O. was supported by the Breakthrough award from the Damon Runyon Cancer Research Foundation (DRG-2378–19), the Dartmouth Innovations Accelerator for Cancer, and a P20 grant (P20GM113132) from the National Institute of General Medical Sciences (NIGMS) of the NIH. R.I.G. was supported by an Outstanding Investigator Award (R35CA232115) from the National Cancer Institute (NCI) of the NIH. We thank the Dartmouth LSE Microscopy Facility for their help with confocal imaging. We thank Kaitlin Taylor for helping with cell-based experiments and Paola Montenegro for helping with mouse brain sections.

## Author contributions

E.A.O., and R.I.G. designed and supervised the research. E.A.O. performed most of the experiments and bioinformatic analyses. I.B., A.T., S.R.J., X.Y., and R.H.A. conducted experiments, analyzed data, and interpreted the results. A.G. provided expert advice on sarcomas and supervised some xenograft experiments. E.A.O. and R.I.G. wrote the paper with input from the other authors.

## Contact for reagent and resource sharing

Further queries and reagent requests may be directed and will be fulfilled by the lead contact, Richard I. Gregory.

## Supplementary Figures

**Figure S1.**
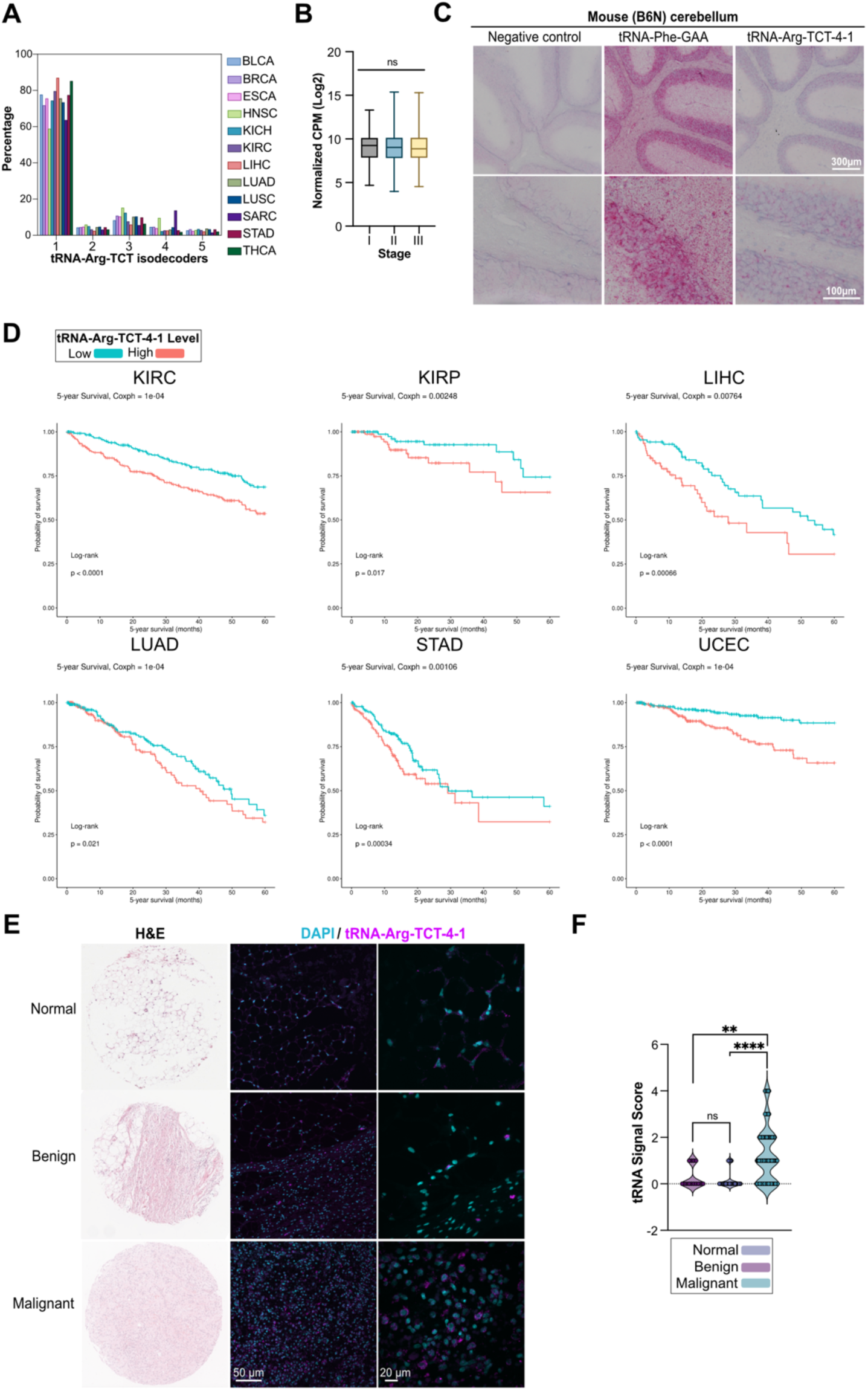
tRNA-Arg-TCT-4-1 expression is associated with malignant phenotypes in cancer and is associated with poor patient outcomes. **A)** tRNA-Arg-TCT isodecoder abundance in multiple human cancers. Normalized data from TCGA. **B)** tRNA-Arg-TCT-4-1 abundance based on the clinical stage of BRCA. **C)** tRNA-Arg-TCT-4-1 (n-Tr20) expression in mice (B6N) cerebellum. **D)** Kaplan-Meier survival curve of multiple cancer types with low versus high tRNA-Arg-TCT-4-1 expression levels. Mean cut-off. Data are from TCGA. Wilcoxon and log-rank test. **E**) tRNA-Arg-TCT-4-1 expression in liposarcoma tumor samples using tRNA-FISH (BaseScope). **F**) Quantification from (C), full TMA. Error bars: Mean ± S.D. P values from Two-way ANOVA with posthoc Bonferroni correction. ns: not significant; ****p < 0.0001.

**Figure S2.**
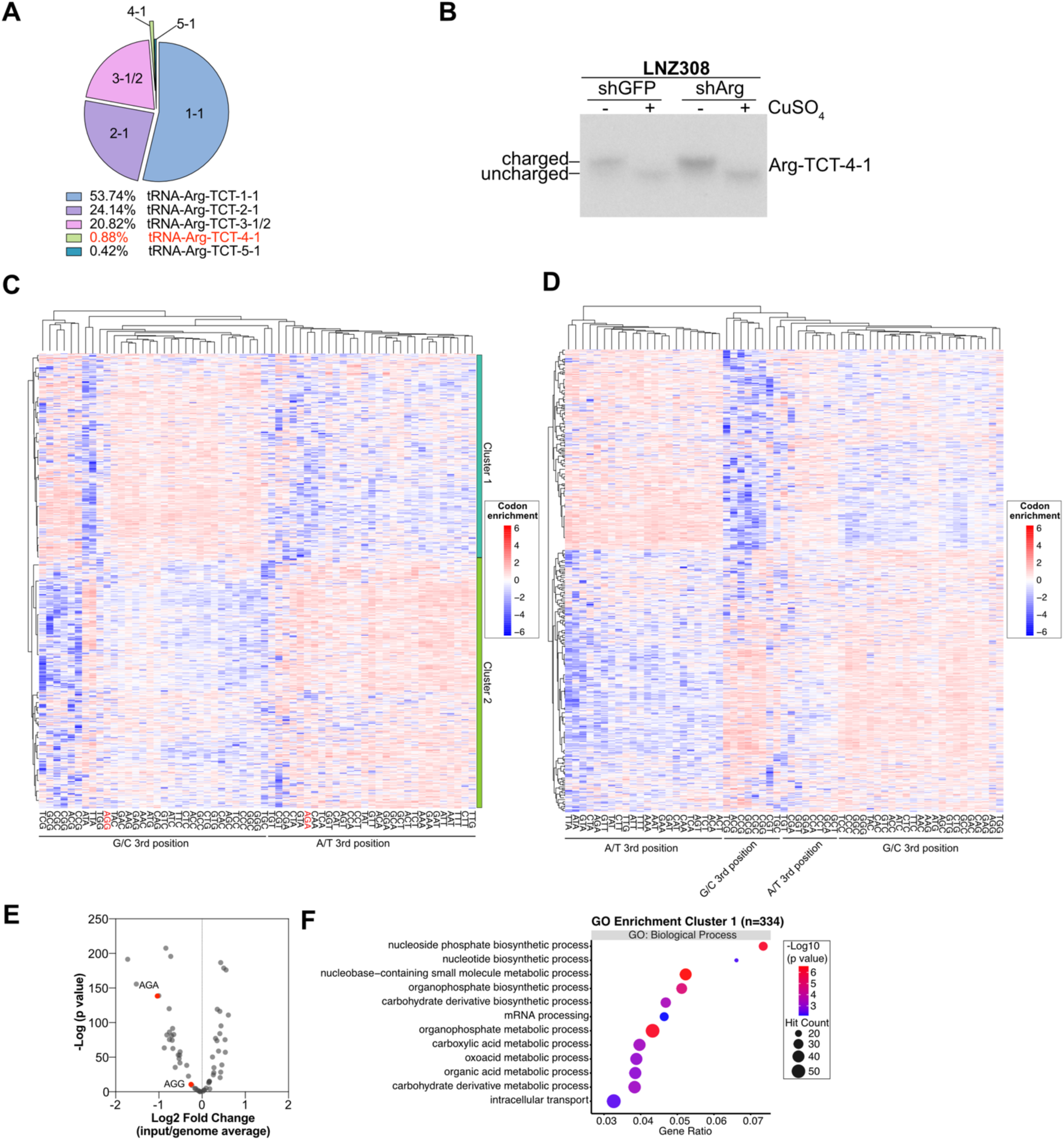
tRNA-Arg-TCT-4-1 inhibition in human glioblastoma cells. **A)** Relative tRNA-Arg-TCT isodecoder contribution to the tRNA-Arg-TCT pool in LNZ308 cells. **B**) Northern blot measuring aminoacylation levels of tRNA-Arg-TCT-4-1 in shArg vs shGFP treated LNZ308 cells. Codon enrichment analyses of downregulated proteins (n=736) (**C**) or a random gene list (n=850) (**D**). Codon enrichment shows the fold change of each codon (61) compared to the genome-wide average. Hierarchical agglomerative clustering, with Euclidean distance and complete linkage. **E)** Codon enrichment (gene input/global genome average) of genes in cluster 1. P-value from Fisher’s Exact Test with Benjamini–Hochberg FDR correction. **F)** Gene Ontology enrichment analysis using downregulated proteins in cluster 1 upon tRNA-Arg-TCT-4-1 inhibition. p<0.05, FC≤1.2.

**Figure S3.**
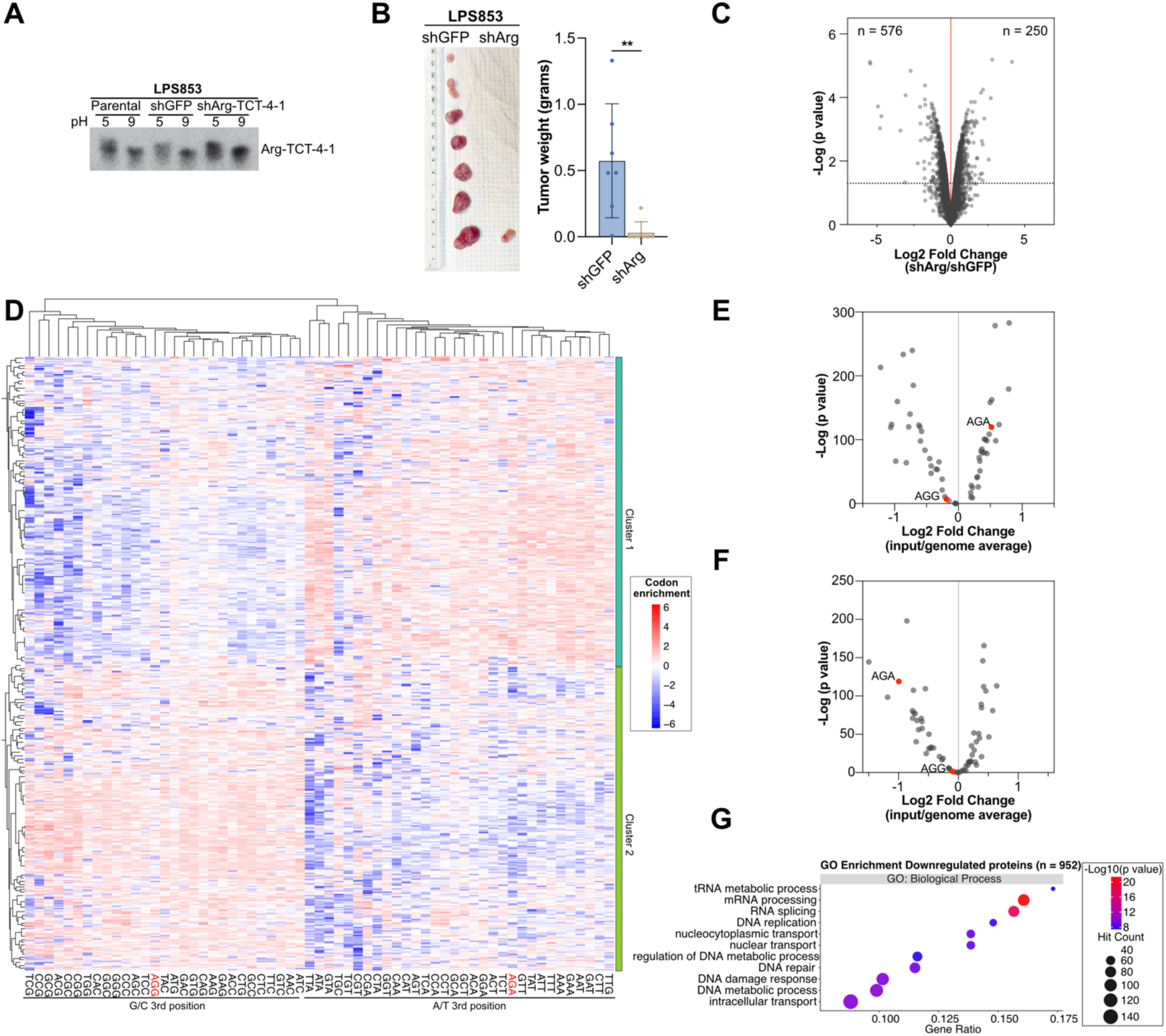
tRNA-Arg-TCT-4-1 inhibition in human liposarcoma cells leads to defects in tumor growth and changes in protein synthesis. **A)** Northern blot measuring aminoacylation levels of tRNA-Arg-TCT-4-1 in shArg vs shGFP treated LPS853 cells. **B)** Quantification of LPS853 tumor weight comparing shArg and shGFP control. Error bars denote mean ± SD. p value from paired Student’s t test. **p < 0.01.**C)** Changes in protein abundance between shArg (heavy) expressing cells and shGFP (light) control cells measured by SILAC-based proteomics; n = 3, moderated t-test. **D)** Codon enrichment analyses of downregulated proteins (n=576). Codon enrichment shows the fold change of each codon (61) compared to the genome-wide average. Hierarchical agglomerative clustering, with Euclidean distance and complete linkage. **E)** Codon enrichment (gene input/global genome average) of genes in cluster 1 and cluster 2 (**F**). P-value from Fisher’s Exact Test with Benjamini–Hochberg FDR correction. **G)** Gene Ontology enrichment analysis using downregulated proteins (n=576) upon tRNA-Arg-TCT-4-1 inhibition. p<0.05, FC≤1.2.

**Figure S4.**
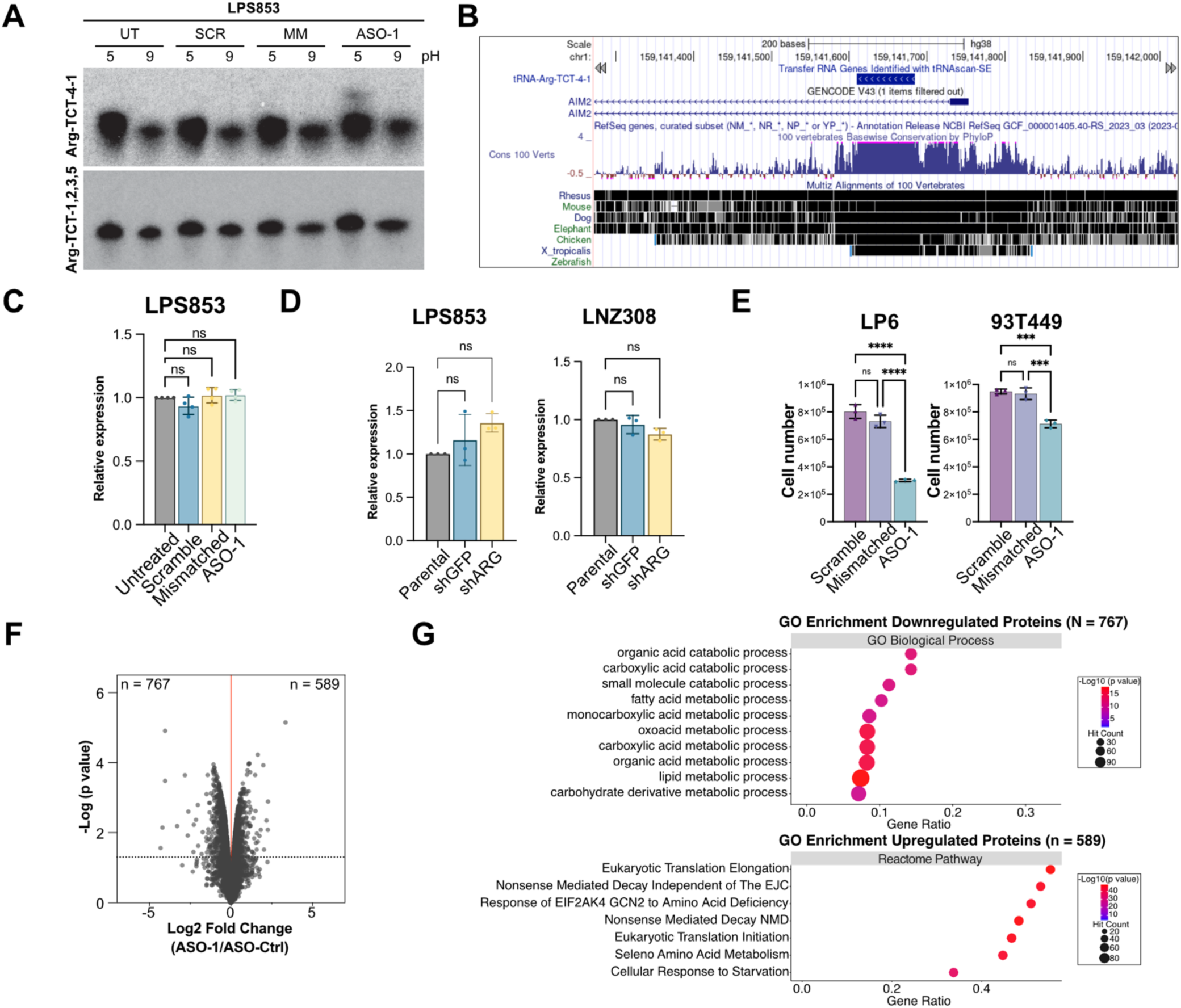
Acute inhibition of tRNA-Arg-TCT-4-1 causes translation defects and decreased cell growth. **A)** Northern blot measuring relative levels of tRNA-Arg-TCT-4-1 in ASO-1 treated LPS853 cells compared to untreated (UT), scramble (SCR), and mismatched (MM) controls. 20 nM, 48 hours post-transfection. **B)** Genomic track of the human tRNA-Arg-TCT-4-1 locus. Captured from UCSC Genome Browser. The expression level of AIM2 in ASO-1 LPS853-treated cells. **(C)** and shArg treatment in LPS853 and LNZ308 cells **(D)**. n = 3 - 4. Error bars: Mean ± S.D. P values from One-way ANOVA with posthoc Bonferroni correction. ns: not significant. **E)** Measurement of cell number upon treatment with ASO-1 in LP6 and 93T449 liposarcoma cell lines. 20nM, 48h after transfection. n = 3. Error bars: Mean ± S.D. P values from One-way ANOVA with posthoc Bonferroni correction. ns: not significant; ***p < 0.001; ****p < 0.0001. **F)** Changes in protein abundance between ASO-1 (heavy) treated cells and ASO-Ctrl (light) treated cells measured by SILAC-based proteomics; n = 3, moderated t-test. **G)** Gene Ontology analysis using Up or Down gene lists from (**F**).

**Figure S5.**
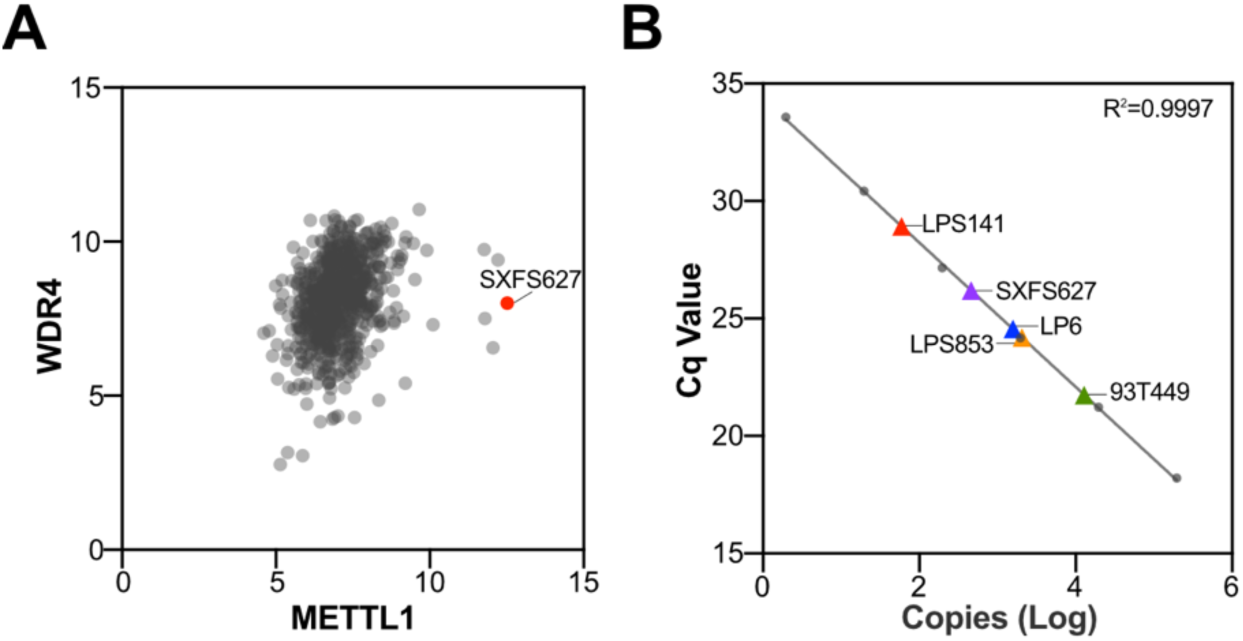
Acute inhibition of tRNA-Arg-TCT-4-1 in a sarcoma PDX. **A)** Normalized METTL1 and WDR4 expression levels in various PDXs. Data from Charles River Laboratories Germany GmbH. **B)** tRNA-Arg-TCT-4-1 expression level in various LPS cell lines and SXFS627. Shown average from technical triplicates.

## Tables

**Table S1.** Data from LNZ308 SILAC Proteomics comparing shArg vs shGFP.

**Table S2.** Ribo-Seq data comparing shArg vs shGFP in LPS853 cells.

**Table S3.** Data from LPS853 SILAC Proteomics comparing shArg vs shGFP.

**Table S4.** Data from LPS853 SILAC Proteomics comparing ASO-1 vs ASO-Ctrl.

**Table S5.** List of oligonucleotides used in this study.

